# HDI-STARR-seq Identifies Functional GH-regulated Sex-Biased Hepatocyte Enhancers Linked to Liver Metabolism and Disease

**DOI:** 10.1101/2025.11.02.686175

**Authors:** Ting-Ya Chang, David J Waxman

**Affiliations:** Departments of Biology and Biomedical Engineering, and Bioinformatics Program Boston University, Boston MA 02215

**Keywords:** Liver sex differences, hepatocyte chromatin accessibility, HDI-STARR-seq, MASLD, epigenomics

## Abstract

Growth hormone (GH) controls sexual dimorphism in hepatocyte gene expression programs governing lipid metabolism, bile acid synthesis and xenobiotic processing, which contribute to sex differences in metabolic dysfunction-associated steatotic liver disease (MASLD) risk. Despite extensive study of GH-regulated sex differences in gene transcription, the functional *cis*-regulatory hepatocyte enhancers that orchestrate these sex-dependent metabolic programs remain largely unknown. Here, we integrated single-nucleus multiomic profiling of hepatocyte chromatin accessibility with *in vivo* functional enhancer assays to identify and validate GH-responsive, sex-biased hepatocyte enhancers in intact mouse liver. We constructed a tiled HDI-STARR-seq library of 23,912 reporters spanning 1,839 liver ATAC regions and delivered it to liver by hydrodynamic injection, enabling functional assessment of enhancer activity *in vivo* across distinct biological conditions. Reporters representing 840 ATAC regions showed sex-biased and/or GH-regulated enhancer activity, in many cases mirroring the regulation of their chromatin accessibility in hepatocytes, validating these sites as functional, physiologically regulated enhancers. The regulated enhancers were enriched for activating histone marks (H3K4me1, H3K27ac), for binding sites for the GH-activated transcriptional regulator STAT5, and for the STAT5-dependent, sex-specific repressors BCL6 and CUX2. Further, *de novo* motif analysis identified binding sites for HNF4A and for several novel factors specifically enriched at the regulated enhancers. Sex-biased and GH-regulated functional enhancers were linked to both MASLD-enabling and MASLD-protective genes, suggesting that GH-dependent chromatin remodeling at these loci contributes to sex-differential metabolic disease susceptibility. This integrated *in vivo* approach defines a validated set of GH-regulated hepatocyte enhancers through which chromatin accessibility and transcription factor binding drive sexual dimorphism in hepatic metabolism and sex-specific MASLD risk.

## Introduction

Growth hormone (GH)–regulated sex differences in hepatic gene expression underlie sex differences in lipid and drug metabolism, xenobiotic clearance and susceptibility to/progression of metabolic liver disease, as seen in both humans and in mouse models (1). GH, acting via its sex steroid-determined, sex-differential temporal patterns of pituitary secretion (2,3), controls the sexually dimorphic expression of several hundred hepatocyte-expressed genes (4) and is a key endocrine regulator of sex-dependent liver function (5,6). GH activates intracellular signaling cascades that lead to distinct, sex-specific expression of cytochrome P450 (*Cyp*) and other metabolic genes, including genes active in lipid metabolism (7,8). Many of these downstream transcriptional outputs are mediated by the TF STAT5 (9–11). Upon binding to GH receptor at the hepatocyte cell surface, GH induces conformational changes leading to activation of JAK2, a GH receptor-linked tyrosine kinase that phosphorylates STAT5, promoting its dimerization (12). Activated STAT5 dimers translocate to the nucleus, where they bind TTCNNNGAA consensus motifs (13) and increase transcription of linked target genes (11). GH-STAT5 signaling plays a pivotal role in diverse physiological processes, including body growth (10,14), cell cycle regulation (15) and hepatic metabolism of lipids, bile acids, steroids and drugs (10,16,17). Dysregulation of GH-activated STAT5 signaling has been implicated in liver disease and its progression from steatosis (7,17,18) to fibrosis and hepatocellular carcinoma (8,19).

Two liver STAT5 isoforms, encoded by the genes *Stat5a* and *Stat5b*, mediate the actions of GH in regulating the transcription of sex-biased genes, with active liver STAT5b protein much more abundant than STAT5a (20). Hepatocyte-specific deletion of *Stat5a* and *Stat5b* in adult male liver (21,22) decreases the hepatic expression of many male-biased genes while de-repressing female-biased genes, consistent with earlier findings in global *Stat5b*-null male mice (9). STAT5b can enhance the transcriptional activity of HNF4A, particularly toward male-biased liver target genes (23,24), many of which are co-dependent on HNF4A for sex-biased expression (25). The transcriptional repressor BCL6, which shows male-biased hepatic expression, is regulated by GH and STAT5 (26) and has a DNA recognition motif that partially overlaps with the STAT5 consensus site, which enables BCL6 to inhibit a subset of liver STAT5 transcriptional responses (13,27). A second GH/STAT5-regulated transcriptional repressor, CUX2, shows female-specific hepatic expression (28) and enforces liver sex differences by repressing many male-biased genes and activating a subset of female-biased genes in female liver (29,30). Together, these findings define a GH-responsive transcriptional network in which STAT5 and HNF4A are major transcriptional activators whose actions are modulated by the sex-biased repressors BCL6 and CUX2 to shape sex-dependent liver gene expression programs.

DNA accessibility plays a crucial role in modulating sex-dependent gene expression by controlling chromatin access for STAT5, BCL6, CUX2 and other liver TFs. Genome-wide DNase-seq and chromatin-state mapping have identified a few thousand sex-biased mouse liver DNase-hypersensitive sites that strongly co-localize with enhancer marks and predict sex-biased gene expression, implicating differential chromatin accessibility as a primary determinant of sex-dependent transcriptional programs (31,32). Plasma GH pulsatility (33) and natural genetic variation (34) both impact the sex-bias of these accessible chromatin regions, dynamically turning accessibility on or off in specific genomic regions in a sex-dependent manner, thereby switching nearby sex-biased genes between male-like and female-like expression states. Further, 3D genome organization studies demonstrate that sex-biased distal enhancers physically contact target gene promoters *via* chromatin looping interactions; and loss of cohesin, which mediates long-range DNA looping, selectively suppresses expression of male-biased genes with distal sex-biased enhancers (35), firmly linking sex-biased chromatin accessibility to sex-biased liver gene expression output. Despite these advances, a critical gap remains: prior epigenomic studies infer regulatory function from correlations between accessibility, histone marks and gene expression, leaving key questions unresolved. First, which of the thousands of sex-biased accessible chromatin regions act as functional, *bona fide* enhancers *in vivo*? Second, does GH signaling and its sex-specific temporal secretion pattern directly modulate liver enhancer activity in the intact endogenous hormonal and metabolic context? Third, are these putative regulatory chromatin sequences linked to genes controlling disease-relevant metabolic pathways implicated in sex-differential MASLD susceptibility?

Liver function is highly dependent on its specialized sinusoidal architecture and cellular heterogeneity, encompassing multiple cell types with distinct phenotypes and functions, including hepatocytes, liver sinusoidal endothelial cells, immune cells and hepatic stellate cells (36–38). Single-cell and spatial transcriptomics studies have revealed pronounced zonation of hepatocyte and non-parenchymal cell gene expression across the liver lobule, as well as substantial intercellular heterogeneity in metabolic pathways and disease-relevant programs (39). Critically, recent single-nucleus (sn)RNA-sequencing analysis established that sex-biased liver gene expression is largely restricted to hepatocytes and is not a feature of liver nonparenchymal cells, with 93% of sex-biased genes showing expression specifically in periportal and pericentral hepatocyte populations (4). Many sex-biased transcripts exhibited sex-dependent zonation within the liver lobule, highlighting an intricate interconnection between liver sexual dimorphism and metabolic zonation at the single-cell level (4). However, most prior work on sex differences in liver gene expression and sex-biased chromatin accessibility has been performed in bulk liver, limiting the ability to distinguish hepatocyte-intrinsic effects from those mediated by liver non-parenchymal cells (NPCs).

To better understand how hepatocyte-specific, zone-resolved gene regulation contributes to liver sex-biased function and metabolic disease risk, and how GH influences these programs, cell-type-specific and liver lobule zone-resolved multiomic approaches were recently employed to jointly profile chromatin accessibility and gene expression using 10x Genomics Multiome ATAC + GEX technology (40). ATAC-seq peaks and RNA expression were simultaneously profiled from the same individual nuclei, enabling discovery of differentially accessible chromatin regions (DARs) as candidates for functional sex-biased and GH-responsive enhancers linked to target gene expression. Here, we build on those findings and present an integrated functional genomics strategy that combines single-nucleus Multiome technology with HDI-STARR-seq (41) to functionally assay and validate sex-biased, GH-responsive enhancers in intact mouse liver. Guided by hepatocyte-resolved chromatin accessibility and gene expression data, we selected ∼1,000 sex-biased and GH-regulated hepatocyte differentially accessible regions (DARs), many linked to sex-biased genes involved in lipid homeostasis and implicated in GH action and its sex-dependent effects on metabolic dysfunction-associated steatotic liver disease (MASLD) (1), together with several hundred non-regulated control ATAC-seq regions. These regions were synthesized as 23,912 tiled reporters and delivered to mouse liver by hydrodynamic injection (HDI) to assay enhancer activity *in vivo* across three biological conditions, under endogenous hormonal and metabolic environments that are absent from conventional cell culture–based reporter assays. This experimental design enabled us to identify physiologically relevant enhancers whose sex-biased, functional enhancer activity mirrors the sex-bias and hormonal regulation of accessibility in hepatocyte chromatin, including enhancers linked to MASLD-enabling and MASLD-protective sex-biased genes, and to discover epigenetic signatures and TF motifs enriched at DARs showing sex-biased, GH-responsive enhancer activity. Together, our findings move beyond correlative epigenomic maps to define a validated set of hepatocyte enhancers through which GH-regulated chromatin accessibility drives sex-dependent function linked to sexual dimorphism in liver metabolism and sex-differential risk of metabolic liver disease.

## Methods

### Animal procedures

All animal experimentation was conducted in accord with accepted standards of humane animal care and in compliance with procedures approved by the Boston University Institutional Animal Care and Use Committee (protocol # PROTO201800698) and ARRIVE 2.0 Essential 10 guidelines (42), including study design, sample size, randomization, experimental animals and procedures, and statistical methods. Male and female CD-1 mice (ICR strain) were purchased from Charles River Laboratories (Wilmington, MA), fed standard rodent chow, and housed in a temperature and humidity-controlled facility on a 12 h light cycle (lights on at 7:30 AM). Reporter plasmids and STARR-seq plasmid libraries were transfected into livers of male and female mice at 7-9 weeks of age by hydrodynamic injection (HDI) on day 0, as described (41). For the cGH-treated male mouse group, mice were implanted on day 1 with an Alzet osmotic minipump set to deliver recombinant rat GH at 25 ng/g of BW/h. For the male vehicle control group, the minipump was loaded with vehicle (30 mM NaHCO_3_) and implanted on day 1. Ten days later, on day 11 after HDI injection, livers were harvested and stored at −80°C. Recombinanat rat GH depleted of bacterial lipopolysaccharide (≤ 7.7 to 15.5 EU/mg) (source: Dr. Arieh Gertler, Protein Laboratories Rehovot, Ltd., Rehovot, Israel) was dissolved in 30 mM NaHCO_3_ and 0.15 M NaCl containing 0.1 mg/ml rat albumin (Sigma-Aldrich, catalog no. A6414) (43). Mice were euthanized under CO_2_ anesthesia by cervical dislocation at 8-10 wk of age and livers were collected at a consistent time of day (between 10:30 AM and 11:30 AM) to minimize the impact of circadian rhythms on gene expression (44). Livers were excised and flash frozen in liquid N_2_ then stored at −80°C.

### In vivo transfection by HDI and validation by luciferase assay

Naked plasmid DNA was delivered to mouse liver by HDI via the tail vein, as described (41). Briefly, 30 μg of pooled STARR-seq plasmid was mixed with 3 μg of plasmid pGYL18 encoding Renilla luciferase and delivered by rapid tail vein injection (HDI) into male and female mice on day 0. The effectiveness of HDI and of plasmid delivery to the liver was verified for each mouse by Renilla luciferase assay, as described (41). Liver extracts were prepared at the same time for each mouse group to minimize batch effects and assayed in technical triplicates. The absolute Renilla activity of each liver was used to assess the success of HDI delivery.

### Single-cell transcriptional and chromatin accessibility profiling of mouse liver

Single cell-based mouse liver gene expression and chromatin accessibility profiles were determined for each biological group using 10X Genomics MultiOme technology, which implements both snRNA-seq and snATAC-seq analysis in the same individual nuclei. The 10X Genomics MultiOmic datasets were analyzed as described (40) using Cell Ranger ARC (v.2.0.0) for initial processing, followed by analysis using Signac (v1.1.0) (45) and Seurat (4.1.0) (46). A total of 127,957 ATAC-seq peaks were identified across all samples and biological groups using the MACS2 peak caller within the Signac package (implemented using the CallPeaks function), which gives ATAC-seq sequence read density data for each biological condition and in each cell type. The CreateChromatinAssay function within Signac was applied to aggregate the genomic ranges of all peaks from the individual sets of liver samples to generate a ‘peaks’ assay.

### Differentially accessible regions (DARs) and differentially expressed genes

Differential chromatin accessibility, and differential gene expression, between biological conditions (male, female and cGH-treated male) was computed (40) using 10x Genomics Loupe Browser 6.0 (https://www.10xgenomics.com/), which uses the exact negative binomial test to test for differences in mean ATAC-seq sequence read density (cut sites per cell) in a given ATAC region, or differences in gene expression between groups of cells. Differential ATAC-seq peaks between cell types were initially identified using the Loupe Browser’s integrated Global Distinguishing function with a differential expression threshold of Log2 (fold change) >1 at FDR (Benjamini-Hochberg) < 0.05. ATAC-seq peaks found in 3% or fewer of nuclei, in the case of non-hepatocyte clusters, or in fewer than 5% of nuclei in the case of hepatocytes were removed. Cell type-specific DARs were identified by excluding peaks found to be cell type-specific DARs for 2 or more clusters. We used the Loupe Browser’s integrated Locally Distinguishing function to calculate condition-specific DARs across five different pairwise comparisons: Male vs. Female-F1, Male vs. Female-F2, Male vs. cGH-treated male (M-cGH), Female-F1 vs. M-cGH, and Female-F2 vs. M-cGH, with a significance threshold of adjusted P-value (Bonferroni-corrected FDR) <0.05, except as noted (and where Female-F1 and Female-F2 represent two independent pools of nuclei from control female mouse livers). We used the same approach, based on the Loupe Browser’s Locally Distinguishing function, described previously (4), to identify differentially expressed genes for the same five comparisons.

### ATAC-seq peak to target gene feature linkages

ATAC-seq peak-to-gene correlations were used to associate sex-biased DARs with their inferred gene targets using feature linkage analysis (40). Briefly, we used the peak-to-gene linkage method embedded in 10x Genomics Cell Ranger ARC (v2.0.1) to link ATAC-seq peaks to specific genes based on Pearson correlations between ATAC-seq peaks and genes, which gives a direct and interpretable measure of the linkage strength (linkage score, values ranging from −1 to 1). We used the set of 127,957 ATAC-seq peaks and 75,798 genes, including some 52,000 mouse lncRNA genes represented by the custom GTF file used in our snRNA-seq analysis (47), to identify 226,841 ATAC-seq peak−gene linkages across the genome, 77% of which involve positive correlations. The subset of these linkages relevant to the ATAC-seq regions we analyzed here is shown in Table S8 and Table S4B. Individual linkages were characterized by the ATAC peak to gene TSS distance, the linkage score (Pearson correlation) and the linkage significance (−log(p-value)).

### ATAC regions selected for STARR-seq reporter library

We selected 1,839 of the 127,957 ATAC-seq peaks regions identified by 10X Genomics MultiOme analysis for HDI-STARR-seq assay of enhancer activity in mouse liver (Table S4A). The ATAC regions selected are in 3 groups: **1)** ATAC regions whose chromatin accessibility is sex-dependent and/or responsive to cGH infusion (Table S2; n= 1,031 ATAC regions, of which 64 regions also belong to the TAD (topologically associating domain)-based group and one is in the pericentral hepatocytes-enriched group), as detailed below; **2)** 749 ATAC regions grouped within 10 specific TAD regions (Table S3A); and **3)** 124 ATAC regions that show significant liver cell type specificity (Table S3B).

### Selection of sex-biased and/or cGH-regulated DARs

A total of 1,031 sex-biased and/or cGH-regulated DARs (Table S2A) were selected for cloning into the STARR-seq library. 612 of these DARs were both sex-biased and cGH-regulated: 360 were male-biased and cGH-repressed, and were linked to 235 male-biased, cGH-repressed genes (Pick1, Table S2B) but were not linked to any genes that were male-biased but not cGH-regulated, or genes that were female-biased genes regardless of GH responsiveness; and 252 were female-biased and cGH-activated DARs linked to 166 female-biased, cGH-induced genes (Pick2, Table S2C) but were not linked to other sex-biased genes (Table S4B, columns W, AD, AK, AR; Table S8). Subsets of both DAR sets also mapped to non-sex-biased genes. 611 of these 612 ATAC regions are found on autosomes (Table S4A). Other sets of DARs included in the STARR-seq library: **1)** 102 male-biased DARs that were stringently not cGH-responsive DARs, 20 of which are on chrY; **2)** 186 female-biased DARs that were stringently not cGH-responsive, 41 of which are on chrX (Table S4A); **3)** 40 male-biased DARs and 20 female-biased DARs that are regulated by cGH but were not significantly linked by the MultiOme analysis to any protein-coding gene or liver-expressed lncRNA gene (47), likely due to their large distance (> 80 kb) from the nearest gene (Table S4E, columns J-K; Pick6 group, Table S2A, Table S4B). These DARs showed HDI-STARR-seq activities similar to those of the sex-biased and cGH-responsive DARs linked to sex-biased genes, including regarding the frequency of regulated reporters (Table S2A); and finally, **4)** 71 sex-biased, cGH-regulated DARs with potential to serve as repressors of sex-biased gene expression (Table S2A, Pick4 and Pick5 DARs). Thus, we selected 62 male-biased and cGH-repressed DARs that were linked to either female-specific and cGH-induced genes (n=34 genes) or to female-specific but GH-independent genes (n=22 genes) (Table S2D); this pattern of GH-regulated chromatin accessibility displays a sex bias and direction of cGH response opposite from that expected for enhancers that regulate expression of female-biased genes, suggesting these DARs may repress their linked female-specific genes. Similarly, we selected 9 female-biased and cGH-activated DARs with repressor potential, i.e., they were linked to male-biased and cGH-repressed genes (n=10 genes) or to male-specific and cGH-unresponsive genes (n=3 genes), displaying the opposite sex response patterns based on their linkage (Table S2E).

### STARR-seq reporter design and synthesis

The 1,839 ATAC regions selected for inclusion in the STARR-seq library were tiled to generate a total of 23,950 STARR-seq reporter sequences, of which only 23,912 reporters were included in the final design (Table S5) owing to issues relating to the liftOver between mouse genome mm10 and mouse genome mm9. Each ATAC region was extended by 70 bp on each end and was tiled using a series of 180 nt reporter sequences, arranged to include 125 nt overlap and a 55 nt shift between neighboring reporters (Fig. 2C). This tiling design minimizes the risk of gaps in reporter coverage of an ATAC region that can arise due to a loss of reporter sequence representation during cloning. Each reporter was synthesized as a 210 nt oligonucleotide using SurePrint HiFi oligo synthesis (Agilent, Inc) and is comprised of the 180 nt reporter sequence flanked on each end by a 15 nt adapter sequence. The pool of 23,912 reporters was then cloned into pPromALB_STARR-seq (plasmid pTYC10; Addgene # 220289) (41), as described below. Each 15 nt adaptor sequence serves as a PCR seed and facilitates the generation of the 210 base pair double-stranded DNA library required for STARR-seq library preparation. Illumina primer sequence #1 is 5’-ACGCTCTTCCGATCT-3’, and Illumina primer sequence #2 is 5’-AGATCGGAAGAGCAC-3’. Sequences of ATAC regions and their mouse genomic coordinates are shown in Table S5.

### STARR-seq plasmid library construction

Each reporter sequence was PCR amplified from the SurePrint HiFi oligo synthesis product using specific PCR primers (Table S1) as described below. We set up 6 individual 25 μl PCR reactions (10 ng of oligo pool, 12.5 μl of 2x KAPA HiFi HotStart master mix (Cat No. KK2601, Roche), 2.5 μl of 5 μM forward primer, 2.5 μl of 5 μM reverse primer per 25 ul reaction) using PCR primers ON8352 and ON8353 (Table S1). PCR reactions were set to generate an HA-adaptor-ligated DNA library under low amplification conditions using the following cycling conditions: 98°C for 45 s, followed by 10 cycles of 98°C for 15 s, 65°C for 30 s, 72°C for 45 s, and final elongation step for 1 min at 72°C. The HA-adaptor-ligated DNA was purified in two rounds of 1x SPRI bead selection and then cloned into pPromALB_STARR-seq vector (Fig. S1). Next, we used a 2:1 molar ratio of insert to 125 ng SalI/AgeI-digested pPromALB_STARR-seq plasmid in two 10 μl In-Fusion® HD Cloning Plus reactions (TaKaRa, cat. #638910) to assemble STARR-seq reporter plasmid libraries via the homology arms. The reaction products were pooled and precipitated using sodium acetate/ethanol, resuspended in 10 μl EB buffer (10 mM Tris HCl, pH 8.5) and stored at −80°C prior to use (48). Two 2.5 μl aliquots of the precipitated material were transformed separately into 20 μl of electrocompetent MegaX DH10B bacterial cells (ThermoFisher Scientific, cat. # EC0113) using Gene Pulser Xcell Total System (Bio-Rad) and Gene Pulser Electroporation Cuvettes (0.1 cm gap, Bio-Rad; cat No. 1652089) under these conditions: 2 kV, 25 µF, 200 ohms. LB bacterial culture medium (1 L) was used to amplify the STARR-seq plasmid library, which was then purified using PureLink™ Endotoxin-Free Maxi Plasmid Purification Kit (ThermoFisher Scientific, cat. #A31231) to obtain ∼ 2 mg of the final STARR-seq plasmid library, STARR-TYC8.

### Preparation of plasmid, transfected DNA and transcribed RNA reporter Illumina sequencing libraries

The following methods, adapted from published STARR-seq protocols (48) and detailed previously (41), were used in a nested PCR amplification protocol to prepare 3 types of Illumina sequencing libraries used to determine the transcriptional enhancer activity of each ATAC region-associated reporter sequence (putative regulatory region) to be assayed by HDI-STARR-seq. 1) A plasmid library was prepared from the same batch of purified plasmid DNA injected into each mouse by HDI. Plasmid DNA was used directly as a PCR template in a nested PCR reaction for library construction (primers ON8381, ON8382; Table S1); 2) a DNA library was prepared from DNA extracted from flash frozen liver tissue collected 10 days after in vivo transfection by HDI (primers ON8381, ON8382); and 3) an RNA library was prepared from the transcribed RNA (reporter sequences) extracted from each HDI-transfected liver (primers ON8379, ON8380). STARR-seq reporter plasmid DNA remaining in the liver 10 days after HDI was extracted using a commercial QIAprep spin miniprep kit (QIAgen, cat. # 27106). The liver tissue (200 mg frozen liver) was initially homogenized in 750 ul of QIAgen Buffer P1 using a rubber-tipped micropestle and passed through a 20G needle three times to homogenize large tissue pieces. The extracted plasmid DNA was bound to a QIAprep spin column, eluted in EB buffer and used as a PCR template in a subsequent nested PCR reaction using primers ON8381 and ON8382. To extract transcribed reporter RNA, 120 μg of total liver RNA was isolated from 60-100 mg liver tissue from each mouse by TRIzol guanidinium thiocyanate-phenol-chloroform extraction followed by polyA selection using oligo(dT) beads (New England Biolabs, cat. #E7490L). The transcribed RNA was digested with Turbo DNase I (Ambion, cat. # AM2238) at 37°C for 30 min, reverse transcribed using primer ON8731 for STARR-seq RNA first strand synthesis and Superscript III first-strand synthesis supermix (Invitrogen, cat. # 18080-400) and treated with 10 μg RNase A per ml (Fermentas, cat. # EN0531) for 1 h at 37°C. The cDNA obtained was purified using 1.8x SPRI beads and then used as a PCR template in the nested PCR reaction.

The nested PCR reaction requires one set of primers specifically targeting STARR-seq reporter DNA or RNA followed by amplification using generalized Illumina dual index primers. Primers ON8381 and ON8382 (Table S1) were designed to amplify the STARR-seq plasmid DNA libraries by specifically targeting the unspliced intron incorporated into the original plasmid sequence and present in the plasmid and DNA libraries but spliced out of the transcribed reporter RNA sequences. Primers ON8379 and ON8380, which straddle the splice junction, were designed to specifically amplify only the transcribed and spliced RNA molecules (41). The second, nested PCR reaction was performed for all three library types (plasmid, DNA and RNA) using NEBNext® Multiplex Oligos for Illumina (NEB cat. # E7600S) to further amplify the material and attach Illumina i5 and i7 barcoded sequences used for Illumina multiplex sequencing, performed at Novogene, Inc. (Sacramento, CA).

### Enhancer identification in TAD regions with sex-biased genes

TADs are fundamental units that compartmentalize genomic features and help constrain interactions of cis-regulatory sequences to their within-TAD genic targets. We examined 6 TAD regions showing a strong tendency toward sex-bias that contain genes encoding important liver metabolic functions, such as cytochromes P450 (CYPs), solute carrier (SLC) family members, cytosolic sulfotransferases (SULTs), UDP glucuronosyltransferases (UGTs), and 3β-hydroxysteroid dehydrogenases (Hsd3b). As controls, we examined ATAC regions from 4 TADs that do not contain any sex-biased genes or sex-biased gene linkages (TAD_203, TAD_1519, TAD_4080, TAD_4241). ATAC regions within the selected TADs were incorporated in the STARR-seq library (Table S3A).

### Liver cell type-specific ATAC regions

The liver MultiOme dataset was used to profile open chromatin regions for each liver cell type or subtype: hepatocytes (including periportal, midlobular and pericentral hepatocytes), liver endothelial cells, hepatic stellate cells (HSC), immune cells, Kupffer cells and cholangiocytes. For each liver cell type (cell cluster), we selected 20 ATAC regions showing the highest specificity for that cell type and defined them as cell-type specific ATAC regions based on the following criteria for the ATAC-seq signal: log2 (fold change) > 1 using the Loupe Browser’s integrated Global Distinguishing function, FDR (Benjamini-Hochberg) < 0.05, and the ATAC-seq signal detected in > 3% of nuclei (> 5% for hepatocytes). ATAC regions that met the criteria for cell type specificity for 2 or more liver cell clusters were excluded. Very few liver cell type-specific ATAC regions were identified for midlobular hepatocytes and cholangiocytes (Table S3B, Table S4C).

### Sequencing analysis of STARR-seq libraries

Paired-end sequence reads, each 150 base pairs long, were generated for the STARR-seq plasmid, DNA, and RNA libraries. A custom pipeline was developed to handle the raw sequencing data (41). This pipeline takes raw FASTQ files as input and produces various quality control metrics, such as FASTQC reports (version 0.0.13.2 of FASTXToolkit), identification and removal of contaminating adapter sequences (using Trim_galore version 0.4.2), and determination of insert-size length distributions for paired-end read sequencing using Picard (version 1.123). The reads were aligned to the mouse genome (version mm9) using Bowtie2 (version 2.2.6) (49). Subsequently, all identified peak sets were filtered to eliminate peaks overlapping ENCODE blacklisted regions (50). An average of 9.5 million fragments were sequenced for the RNA libraries and 27.1 million fragments for the DNA and plasmid libraries (Table S6). Almost all (96-99%) of the plasmid, DNA and RNA library reads mapped to the 23,912 predesigned reporters for all samples except for library G202M16, a negative control for HDI in the absence of a STARR-seq reporter library. Five RNA sequencing samples were excluded from the downstream analysis (G202M1, G202M2, G202M6, G202M10, G202M12) due to low Pearson’s correlations with other RNA libraries from the same biological condition (Table S6). Similarly, DNA sequencing sample G202M18 was removed due to its low Pearson’s correlation with the other 3 DNA libraries and the 2 plasmid libraries (Table S6). All Raw fastq sequencing files are available at GEO (https://www.ncbi.nlm.nih.gov/geo/) under accession number **GSE315771**.

### STARR-seq library characterization

STARR-seq plasmid sequencing reads, and liver-extracted STARR-seq DNA library sequencing reads, were mapped to the set of 23,912 reporters. Since adjacent reporter sequences overlap by 125 bp of sequence, we used Bedtools (v2.31.0) to set > 95% overlap (>171/180 nt) as the threshold for assigning a sequence read to a given reporter to avoid counting partial overlaps from neighboring reporters. Sequence reads obtained for each plasmid library mapped with high frequency (98.8%) to the 23,912 designated reporters (Table S6), indicating very low non-specific PCR amplification and minimal sequencing errors. The two technical replicate plasmid libraries were highly correlated based on normalized counts across the 23,912 reporter regions (R^2^=0.984, Fig. S2A). Almost all (>98%) of the inserts were 180 nt long, showing that intact oligos were cloned into the library. The median number of normalized plasmid sequence reads in each reporter region was 322, and 81.9% of the 23,912 reporters were represented within 6.4x of the median frequency (Fig. S2B). The GC content of the 23,912 reporters included in our library design ranged from 19-87%, with the highest frequency observed for reporters at ∼45-50% GC content (Fig. S2C). The PCR-amplified plasmid and liver-extracted DNA libraries average sequence read counts both peaked at ∼55-70% GC content (Fig. S2D), consistent with the known bias against both GC-rich and AT-rich in sequencing libraries prepared using a PCR-amplification step (51), suggesting that differences in reporter representation across the library is largely determined by the sequence of each reporter.

### Qualification of STARR-seq library reporters for downstream analysis

A total of 19,136 STARR-seq reporters (Table S7A, column AU) were qualified for inclusion in downstream analysis based on the following stringent threshold for robust representation in the cloned and bacterially amplified STARR-seq plasmid library, STARR-TYC8: > 40 mapped Illumina sequence read pairs in each of the 3 liver-extracted STARR-seq DNA libraries and in both individual plasmid libraries (Table S7A; Fig. S3B, first 2 columns). 4,776 other reporters did not meet this threshold for consistent STARR-seq library representation and were excluded from downstream analysis. A very similar list of qualified reporters was obtained when the threshold of > 40 mapped Illumina sequence read pairs was normalized to the total sequence depth of each plasmid or DNA library.

### Determination of HDI-STARR-seq enhancer activity

Illumina sequencing libraries prepared from RNA extracted from each HDI-STARR-seq-transfected liver (‘RNA library’) were analyzed using Bedtools to count the number of RNA sequence reads that overlapped each of the 19,136 qualified reporters. Qualified reporters with > 30 mapped Illumina RNA sequence read pairs in all 3 biological replicates livers from untreated male mice, in both biological replicate livers from cGH-treated male mice, <u>or</u> in both biological replicates from untreated female mice were deemed active enhancers. A total of 11,996 (63%) of the 19,136 qualified STARR-seq reporters were thus determined to be active, with values ranging from n=7,099-8,922 for each biological condition (Fig. S3E). The remaining 7,140 reporters (37%) were inactive under all three biological conditions. Further, a total of 4,126 of the 19,136 qualified STARR-seq reporters met these criteria for active enhancers under all 3 biological conditions and were designated robust active enhancers (Table S7, column AT). The robust active reporters mapped to 1,292 ATAC regions (Fig. S3F) were somewhat more highly represented in the STARR-seq plasmid library than the qualified reporters that showed activity under only 1 or 2 of the 3 biological conditions (Fig. S4A), suggesting a higher false positive rate for the latter sets of non-robust STARR-seq enhancers owing to noise associated with genomic regions with lower representation in the STARR-seq plasmid library. We therefore limited our analysis of differences in enhancer activity between biological conditions (see below) to the set of 4,126 robust active reporters. 982 of the 19,136 qualified reporters were designated stringently inactive reporters based on these criteria: ≤ 30 mapped Illumina sequence read pairs in each RNA library across all 3 biological conditions (i.e., in all 7 RNA libraries included in our analysis; Table S7, column AT). Since many of these inactive reporters were poorly represented in the STARR-seq library, we subdivided the 982 stringently inactive reporters into two sets based on mean plasmid library read counts: stringently inactive-High (top 400 reporters), which were used as a background set for downstream enrichment analyses; and stringently inactive-Low (bottom 582 reporters), which were excluded from the analysis (Fig. S4A). A further 6,188 qualified reporters were designated ‘Other’ and were excluded from further analysis. The ‘Other’ designation was assigned based on inconsistent STARR-seq enhancer activity across biological replicates and may relate to low representation of these reporters in the plasmid library (Fig. S4A). The lists of stringently inactive and robust active reporters were highly similar, or nearly identical, respectively when the thresholds for minimum numbers of raw mapped read counts stated above were normalized for the total sequence depth of each RNA library (e.g., 4,125 of the 4,126 robust active reporters were retained when using those alternative criteria).

### Relationship between sex bias and cGH responsiveness of STARR-seq enhancer activity

A linear regression model (52) was implemented in R Studio to assess the relationship between response to cGH treatment (log2 ratio of STARR-seq enhancer activity in cGH-treated male liver to STARR-seq enhancer activity in male liver) and sex bias (log2 ratio of STARR-seq enhancer activity in female liver to STARR-seq enhancer activity in male liver).

### Regulated active STARR-seq enhancers

Robust active enhancers with > 2-fold differences between biological conditions in their STARR-seq reporter activity (defined as mean RNA library sequence reads for a given HDI-STARR-seq reporter, averaged across biological replicates, normalized for total mapped sequence reads) were designated regulated active enhancers (Table S7A). Regulated active enhancers whose pattern of HDI-STARR-seq enhancer activity regulation matched their pattern of ATAC-seq activity regulation were designated positively acting regulated enhancers. One example is the set of DARs whose ATAC-seq activity was both male-biased and cGH-suppressed (Pick1 DAR set, Table S2) and whose RNA reporter(s) showed >2-fold increase in STARR-seq enhancer activity in male compared to female liver as well as a >2-fold decrease in HDI-STARR-seq enhancer activity in cGH-treated male liver compared to control male liver. Another example is the set of DARs whose ATAC-seq activity was male-biased but not cGH-regulated (Pick3_M DAR set, Table S2) and whose reporter(s) showed >2-fold increase in STARR-seq enhancer activity in male compared to female liver but did not show >2-fold decrease following cGH infusion. In contrast, male-biased and cGH-suppressed DARs whose STARR-seq reporter activity showed < 2-fold male-bias and < 2-fold response to cGH infusion were non-regulated enhancer DARs. We also identified a set of male-biased, cGH-repressed DARs linked to female-biased genes (putative repressors; Pick4 DAR set, Table S2), whose reporters showed >2-fold increase in STARR-seq enhancer activity in male compared to female liver and also >2-fold decrease in STARR-seq enhancer activity in cGH-treated male compared to control male liver, and correspondingly for female-biased, cGH-activated DARs linked to male-biased genes (Pick4 and Pick5 DAR sets; Table S2). Finally, 452 robust active enhancers whose reporters showed no regulated STARR-seq enhancer activity in any comparison (Table S9A, column AI) were identified as non-regulated DARs.

### Chromatin state analysis of STARR-seq regions in liver

Chromatin state maps, previously defined for adult male mouse liver, and separately, for adult female mouse liver (32) using ChromHMM (53), were used to determine the enrichment of individual chromatin states in various sets ATAC regions. These chromatin states are based on a 14-state model (states E1-E14) and were previously used to characterize sex differences in chromatin state and their relationships to sex-biased gene expression (32). Bedtools overlap was used to assign the chromatin state determined for each sex to each ATAC region. An ATAC region that overlapped two or more different chromatin states was assigned to the state with the greatest overlap in bp. The distribution of chromatin state enrichments (overall chromatin landscape) was determined separately for male and for female liver for each set of functionally active enhancer region DARs in comparison to a background set comprised of 317 DARs whose reporters showed no regulated STARR-seq enhancer activity in any comparison (i.e., 452 DARs mentioned above, after excluding 82 non-regulated DARs from the Pick1 DAR set and 53 non-regulated DARs from the Pick2 DAR set) (Table S9A column AJ).

### Motif analysis

*De novo* motif discovery was carried out using MEME-CHIP (54) (https://meme-suite.org) run in the classic mode to discover motifs enriched relative to a random model based on frequencies of each nt in the input sequences (Table S13A). MEME-CHIP was also run in the discriminative (non-classic) mode comparing an input set of sequences to a background set of sequences, as specified in the text, using default settings for both modes (Table S13B). For each set of regions analyzed (Table S9A, columns U-X) sequences were input using the coordinates for each region shown in Table S9A. For example, STARR-seq analysis of a set of 360 male-biased and cGH-repressed DARs identified 41 functionally active regulated ATAC regions (i.e., regulated Pick1). Sequences of those 41 ATAC regions were obtained from the mouse mm10 genome using the bedtools getfasta command and used as input. Specified background sets include: the set of 65 stringently inactive ATAC regions (Table S9A, column AK), defined as ATAC regions that contain 1 or more stringently inactive STARR-seq_High reporters (Table S7A, column AT) but do not include any of the 4,126 robust active reporters; and the set of 82 active but not regulated, male-biased, cGH-repressed DARs and the set of 53 active but not regulated, female-biased, cGH-activated DARs (Table S9A, columns V, X). The *de novo* discovered motifs were compared with published mouse TF motifs determined by protein-binding microarrays and obtained from the UniPROBE motif database (55). Separately, Find Individual Motif Occurrences (FIMO) analysis was conducted to identify Stat5a/Stat5b, BCL6, CUX2 and HNF6 motif occurrences within sets of regions by scanning the motif for STAT5A/STAT5B (JASPAR motif ID: MA0519.1), BCL6 (Hocomoco mouse motif ID: BCL6_MOUSE.H11MO.0.A), CUX2 (Hocomoco mouse motif ID: CUX2_MOUSE.H11MO.0.C) and HNF6 (Hocomoco mouse motif ID: HNF6_MOUSE.H11MO.0.A) (56,57) (Table S13C, columns AP-BM).

## Results

### Candidate sex-biased and GH-responsive regulatory regions identified by snATAC-seq + snRNA-seq analysis

We implemented 10X Genomics MultiOme technology (snATAC-seq in combination with snRNA-seq) to identify open chromatin regions that show sex-dependent and/or GH-regulated accessibility in mouse liver nulcei from male, female, and cGH-infused male mice. 46,188 nuclei across the 3 biological conditions passed quality control filters for both the snATAC-seq and snRNA-seq modalities. ATAC-seq data quality was confirmed by TSS enrichment scores of 5.5–6.4. Five major liver cell types were resolved by UMAP and identified using established marker genes (Fig. 1A). 127,957 ATAC regions were identified across all liver cells and characterized regarding sex-bias and the impact of cGH infusion on chromatin accessibility. We linked individual ATAC regions to predicted gene targets using correlations between chromatin accessibility and gene expression; these ‘co-expression’ relationships indicate a shared regulatory mechanism and infer enhancer–gene targeting relationships. For example, a feamle-biased, cGH-activated DAR located 46.1 kb downstream of the *Cux2* TSS (ATAC_39124) showed linkages to 4 nearby genes (Fig. 1B, Fig. 1C): two positive linkages, to female-biased genes (*Cux2*, *lnc4593*), indicate positive regulation, and one negative linkage, to *P2rx4*, indicates negative regulation by the ATAC region (Fig. 1D). The linkage to *Cux2* showed the highest correlation (*r* = 0.41) and significance (–log(*P)* FDR = 70.6).

**Fig 1.**
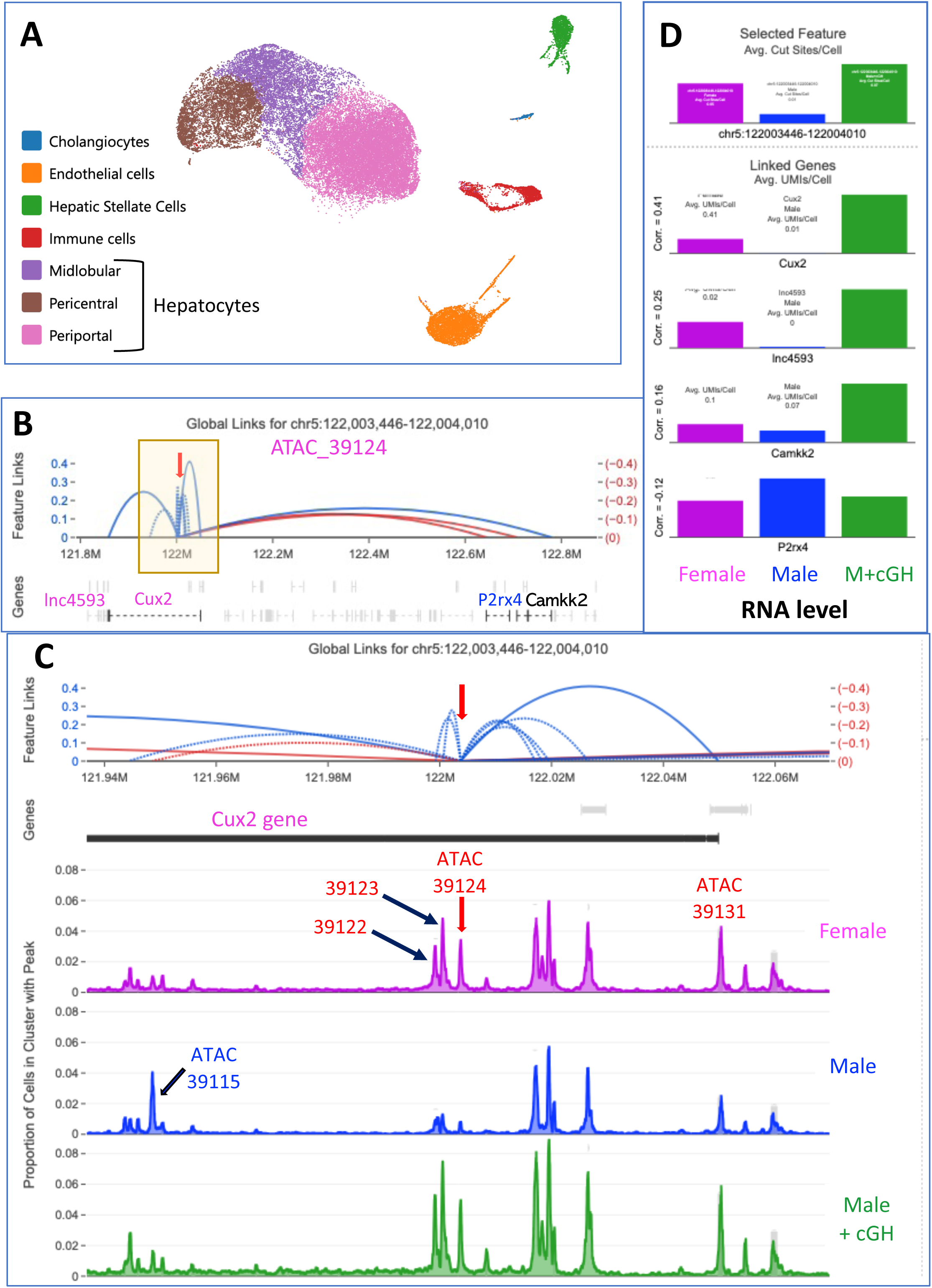
Joint transcriptional and chromatin accessibility profiling of mouse liver. **(A)** UMAP of 46,188 mouse liver nuclei based on 10X Genomic MultiOmic joint transcriptomic + chromatin accessibility profiles harmonized across three biological conditions: male liver, female liver and cGH-treated male liver. Cell types were identified using established marker genes. **(B)** snATAC-seq linkage analysis for female-biased, cGH-activated ATAC_39124 (red arrow, mouse mm10 genomic coordinates indicated), revealing inferred short and long distance positive regulatory linkages (blue arcs) to female-biased genes *Cux2* and *lnc4593*, and negative regulatory linkages (red arcs) to male-biased gene *P2rx4*. Region within yellow box is expanded in panel C. **(C)** Closer view of ATAC_39124 region with linkages (*top*) and normalized ATAC peaks for each biological condition (*bottom*). Select sex-biased and cGH-responsive ATAC peaks are numbered, including negative regulation to male-biased, cGH-repressed ATAC_39115. **(D)** Bar graphs showing ATAC activity (cut sites/cell) for region ATAC_39124 (*top*), and RNA expression levels for the 4 indicated genes in female, male and in cGH-infused male liver (*bottom*), whose linkages to ATAC_39124 are all significant at −log10(FDR) > 5 (Table S4B).

### Selection of DARs for HDI-STARR-seq library

We selected ATAC regions for HDI-STARR-seq analysis, focusing on genomic regions showing cGH-regulated sex differences in accessibility in hepatocytes, the primary liver cell type showing gene expression and chromatin accessibility sex differences (4,40). Three major types of DAR sets were chosen: 1) 360 DARs that were male-biased and cGH-repressed and linked to correspondingly regulated male-biased genes (Fig. 2B, Table S2A, ‘Pick1’), and 252 DARs that were female-biased and cGH-activated and linked to correspondingly regulated female-biased genes (Fig. 2B, Table S2A, ‘Pick2’); 2) 102 male-biased and 186 female-biased DARs whose chromatin accessibility was unresponsive to cGH, subsets on which were on chrY and chrX, respectively (Table S2C, ‘Pick3’); 3) 749 ATAC regions from 10 TAD regions (58) encompassing sex-biased genes important for hepatic lipid metabolism and fatty liver disease, including cytochromes P450 (CYPs), solute carrier (SLC) family members, sulfotransferases (SULTs), UDP glucuronosyltransferases (UGTs), and 3β-hydroxysteroid dehydrogenases (HSD3B family) (Table S3A). Other ATAC regions evaluated include putative repressor sequences (Pick4, Pick5), which are male-biased or female-biased DARs whose regulatory orientation relative to their linked genes suggests repressive rather than activating functions, and sex/cGH-regulated ATAC regions not linked to any genes (Pick6), which may represent long-range enhancers, given their large distances (> 80 kb) from nearest genes (Table S2, Table S4E). These DAR sets all showed similar ranges of HDI-STARR-seq enhancer activity (Fig. S5). Additionally, 124 ATAC regions were included to determine the liver cell type-specificity of HDI-STARR-seq (Table S3B). Overall, we profiled 1,839 ATAC regions (mean 634 nt), tiled with overlapping reporters spaced 55 bp apart to finely map active sub-sequences (Fig. 2C).

**Fig 2.**
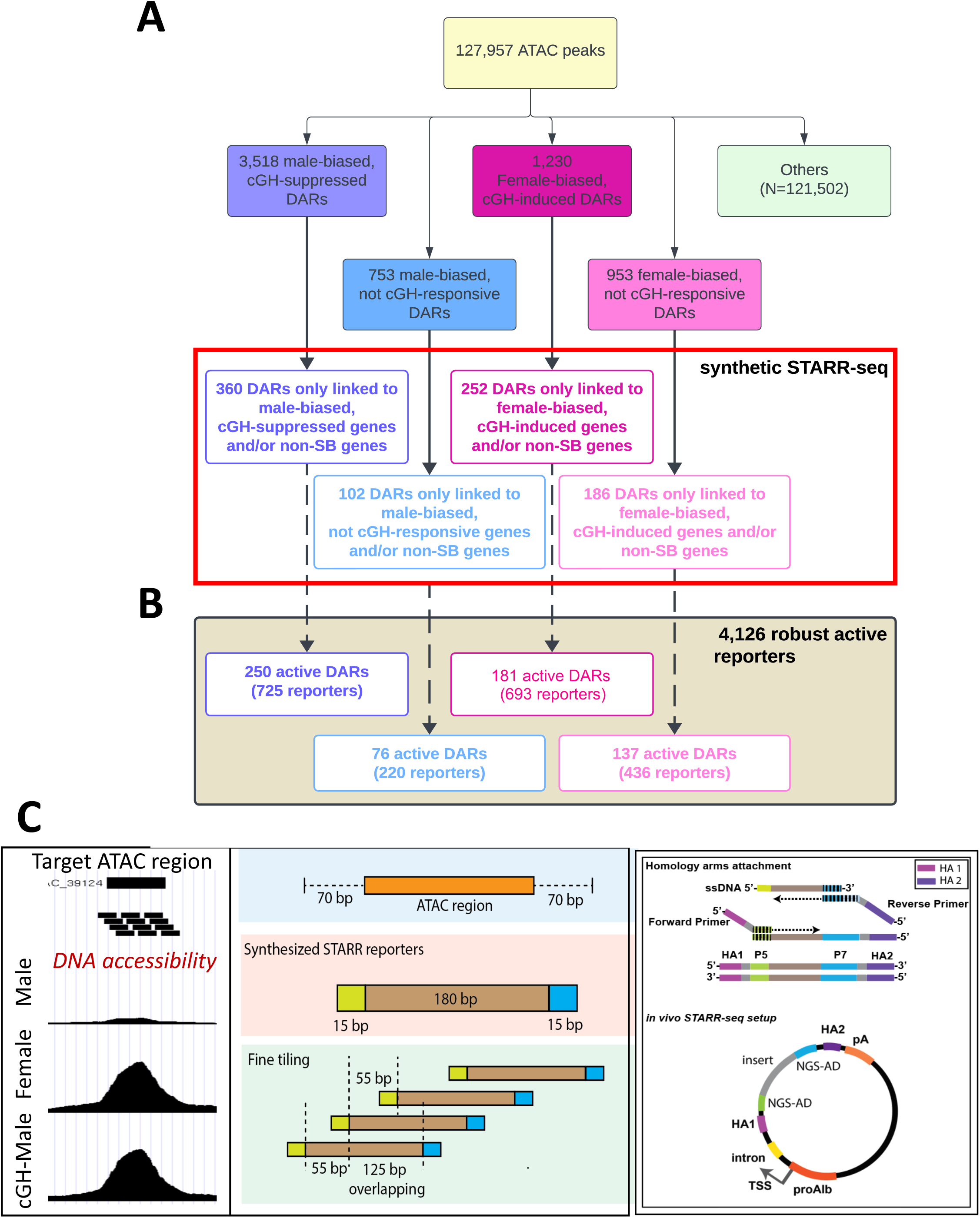
Synthetic STARR-seq reporter library. **(A)** Sex-biased and/or cGH-regulated DARs selected for library (red box) based on the sex bias (SB) and cGH-responsiveness of their ATAC activity and that of their linked genes (Table S2). DARs selected were derived from the full sets of ATAC-seq peaks indicated at top. **(B)** Number of active DARs from each set in panel A with at least 1 robust active reporter. Numbers in parentheses indicate the number of robust active reporters from the full set of 4,126 robust active reporters. For example, 250 of the 360 male-biased, cGH-repressed DARs include a total of 725 robust active reporters from the full set of 4,126. **(C)** Design of synthetic STARR-seq library reporters based on liver ATAC region plus 70 nt of flanking sequences. *Left*, Schematic showing a female-biased ATAC region activated by cGH infusion and a series of tiled reporters. *Middle,* Each reporter is comprised of a genomic sequence flanked by 15 nt of adaptor sequences (olive, blue). Neighboring reporters are shifted by 55 bp and overlap by 125 bp. *Right*, adaptor sequences for PCR, where the primers anneal and incorporate homology arms for the HiFi cloning step. PCR amplification is followed by HiFi cloning to attach homology arms to single stranded DNA oligos (see Methods).

### Active enhancers identified by HDI-STARR-seq

A pool of 23,912 unique single-stranded oligonucleotides (Table S5, Table S7A) tiled across the 1,839 ATAC regions was cloned in bulk into the STARR-seq reporter vector pPromALB_STARRseq (41). The library was high-quality, with 19,136 (80%) of the reporters passing stringent thresholds for robust representation (see Methods). We used HDI to transfect the STARR-seq reporter library into mouse liver and assay the enhancer activity of each ATAC region under 3 biological conditions: male, female, and 10-day cGH-infused male liver. Livers were harvested after 11 days and transfection efficiency was assessed with a co-transfected Renilla luciferase reporter. Low Renilla activity livers, indicating poor transfection, were excluded from downstream analysis (Fig. 3A, asterisks). DNA reporter sequences were recovered from the livers without bias (Fig. S6). RNA reads from the 7 livers showing robust HDI transfection (n=3 males, n=2 females, n=2 cGH-treated males) were mapped to the 23,912 reporters with high efficiency (>98%, Table S6) and general overall consistency within groups (Fig. 3B). In total, 11,996 of the 19,136 reporters (63%) showed enhancer activity under one or more biological conditions and were designated *active enhancers* (Fig. S3E; Table S7A, column AU); 4,126 reporters (21.6%), representing 1,292 ATAC regions, were active under all 3 conditions and are termed *robust active enhancers* (Fig. S3F; Table S7A, column AT).

**Fig. 3.**
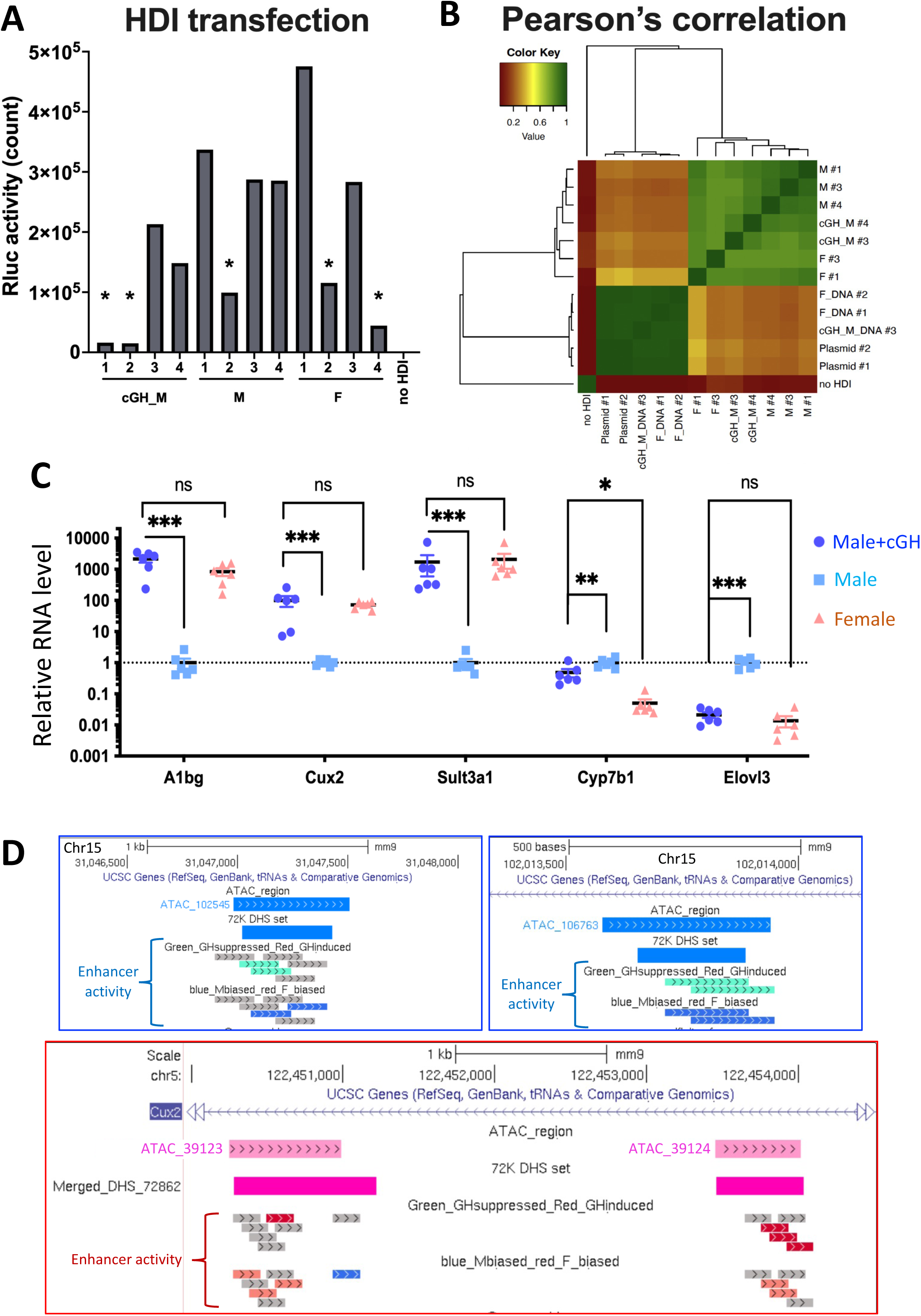
In vivo evaluation of HDI-STARR-seq library transfection and cGH response. **(A)** Validation of reporter plasmid transfection based on Renilla luciferase reporter activity in each of 12 HDI-transfected mouse livers. Five livers with lower luciferase activity (asterisks) were excluded from downstream analyses. **(B)** Clustering of normalized mapped read counts in ATAC regions from biological replicates for 2 plasmid libraries, 3 DNA libraries and 7 RNA (enhancer activity) reporter libraries, based on Pearson’s correlation. DNA and RNA libraries were extracted from HDI livers for each biological condition: M, male; F, female; and cGH-treated male. **(C)** Impact of cGH infusion on RNA levels of the indicated sex-biased genes. Significance: Kolmogorov-Smirnov test, followed by multiple comparison with FDR for multiple comparison correction. Data are mean +/− SEM (n=6 livers/group), with male livers (vehicle pump) mean value = 1. **(D)** UCSC Browser screen shots with examples of sex-biased, cGH-regulated ATAC regions from Pick1 set (*top*) and Pick2 set (*bottom*) whose HDI-STARR-seq enhancer activity shows matched regulation for cGH response (Repressed, green; Induced, red) and sex (male-biased, blue; Female-biased, light red) across multiple concordant reporters (*lower two tracks*). Boundaries of the ATAC region identified in the current study and of the overlapping sex-biased DNase-hypersensitive sites (DHS) identified in bulk liver (‘72K DHS set’) (31) are shown. Female-biased ATAC regions shown within *Cux2* intron are detailed in Fig. 1C.

### Impact of cGH infusion on functional activity of active sex-biased enhancers

Subcutaneous infusion of GH in male mice using osmotic minipumps (cGH infusion) mimics the female plasma GH pattern and overrides the male pulsatile pattern of pituitary GH release, substantially feminizing sex-biased gene expression in mouse liver within 7–10 days (43,59). We used cGH infusion to identify ATAC regions whose enhancer activity responds to changes in temporal plasma GH patterns. cGH infusion for 10 days feminized the liver, confirmed by increased expression of female-biased genes (*A1bg*, *Cux2*, *Sult3a1*) and decreased male-biased gene expression (complete repression, *Elovl3*; partial repression, *Cyp7b1*) (Fig. 3C). Liver stress gene expression, which in a prior study was elevated for up to 4 days following surgery to implant osmotic minipumps (43), was not was observed in response to HDI or 10 days after minipump implantation (Fig. S7). Therefore, the 10-day post-HDI interval is sufficient for recovery from any stress-related responses.

We examined the set of 4,126 robust active reporters to identify those showing sex-dependent or cGH-responsive activity. Of these, 1,397 reporters exhibited HDI-STARR-seq enhancer activity that was either male-biased (n=310), female-biased (n=541), cGH-induced (n=398), and/or cGH-repressed (n=419) (Table S7A, column AW). Concordant effects across neighboring reporters support true local regulation (Fig. 3D). The 1,397 reporters mapped to 840 distinct ATAC DAR regions, including 168 male-biased DARs whose accessibility was repressed by cGH, and 137 female-biased DARs whose accessibility was increased by cGH. Further, our analysis indicated that the sex-bias of enhancer activity was the key determinant of cGH responsiveness. Thus, 87 male-biased reporters showed decreased enhancer activity following cGH infusion, versus only 1 showed an increase (Fig. 4A). Moreover, among the female-biased reporters, cGH infusion increased enhancer activity for 179 reporters, but decreased it for only 4 (Fig. 4A). This pattern mirrors the well-established feminization of liver chromatin accessibility and gene expression following cGH infusion (43,59), supporting the conclusion that HDI-delivered STARR-seq reporters recapitulate physiological GH-regulated enhancer responses.

**Fig. 4.**
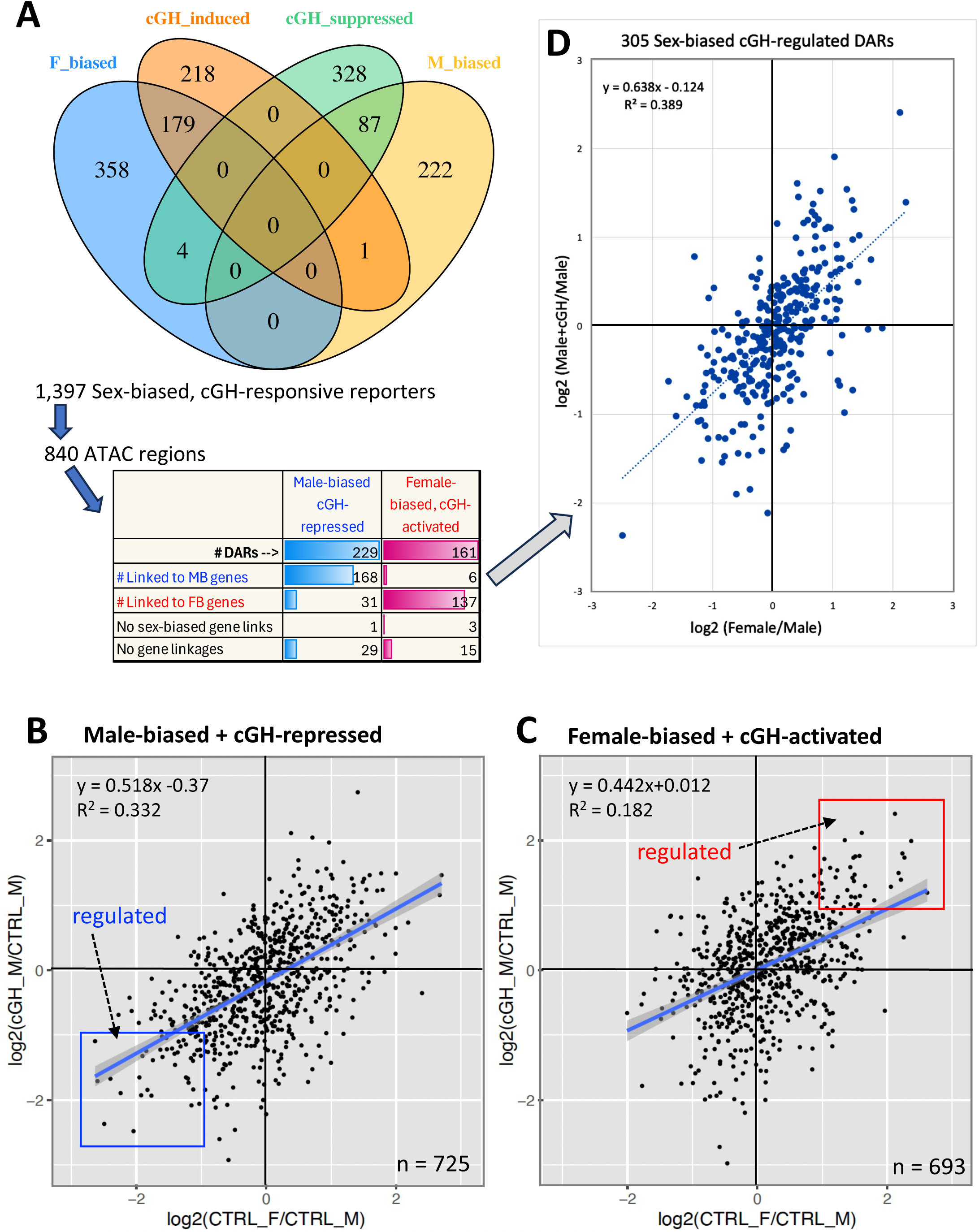
Relationship between sex bias and cGH-responsiveness of HDI-STARR-seq enhancer activity. **(A)** Venn diagram showing overlaps between the 1,397 HDI-STARR-seq reporters whose enhancer activity is sex-biased and/or cGH-regulated (Table S7A, columns AW-BB). These reporters map to 840 DARs, including 229 male-biased (MB) DARs repressed by cGH and 161 female-biased (FB) DARs activate by cGH, whose distributions of gene linkages to sex-biased and other genes are indicated in the table. **(B, C)** Correlation between sex bias of reporter activity and response of reporter activity to cGH infusion in male liver for the set of n=725 robust active reporters (subset of 4,126 such reporters; Table S7A) that map to male-biased, cGH-repressed DARs (panel B, Pick1 DARs) and for n=693 robust active reporters that map to female-biased, cGH-activated DARs (panel C, Pick2 DARs). Blue and Red boxes, reporters whose enhancer activity is regulated by both sex and cGH. Red box: 41 male-biased and cGH-repressed reporters; red box, 43 female-biased and cGH-activated reporters. See Table S7B for other correlation data. Blue lines in panels B, C: best fit linear regression models, with the 95% confidence interval show in gray. **(D)** Mean DAR-level analysis of correlations, as in B and C, applied to 305 DARs: 168 male-biased and cGH-repressed DARs, and 137 female-biased and cGH-activated DAR. Data are mean values of log2 ratios calculated across all robust active reporters mapping to each DAR.

Plotting the fold-change in enhancer activity following cGH infusion versus the sex-bias in enhancer activity revealed strong positive correlations for HDI-STARR-seq reporters mapping to male-biased/cGH-repressed DARs (360 DARs, n=725 reporters) and to female-biased/cGH-activated DARs (252 DARs, n=693 reporters), respectively (Fig. 4B, Fig. 4C). Non-regulated DAR sets showed weaker, less significant correlations (Table S7B, Fig. S8). Separating sex-biased but cGH-unresponsive reporters into autosomal and sex chromosome subsets revealed stronger correlations for autosomal reporters (p = 5.6E–31 vs p = 3.5E–03) (Fig. S8C-S8D), consistent with sex chromosome DARs being regulated by GH-independent mechanisms (40). Importantly, analysis of the reporter datasets averaged at the ATAC region level revealed an even stronger correlation between enhancer activity sex bias and cGH-responsiveness (Fig. 4D).

### Enrichment of regulated active enhancers for DARs with matched response

Enrichment analysis confirmed the close association between the sex bias and cGH responsiveness of HDI-STARR-seq reporter activity and the sex-bias and cGH regulation of endogenous mouse liver chromatin accessibility (Table S7C). Male-biased and cGH-repressed DARs were significantly enriched for male-biased, cGH-repressed enhancer activity compared with female-biased and cGH-activated DARs (ES = 8.3, p = 3.7E-08, Fisher’s exact test). Enrichments for other regulated DAR sets appear in Table S7C. The sex-bias in chromatin accessibility in liver did not differ significantly between the regulated and non-regulated DAR sets (Table S7E). Furthermore, in male liver, no significant difference in chromatin accessibility was found for regulated vs. non-regulated male-biased and cGH-repressed DARs, but in female liver, the regulated female-biased and cGH-activated DARs showed significantly greater accessibility than the non-regulated DAR set (Table S7E). Gene linkage metrics did not differ between the regulated and non-regulated sets of sex-biased, cGH-responsive HDI-STARR-seq enhancers, which showed no differences in the proportion of non-sex-biased genes, distance to linked genes, or linkage significance for either DAR type (Fig. S9, Table S12). Finally, many sex-biased and cGH-regulated DARs were linked to both sex-biased and non-sex-biased genes (Table S8), but the linkages to non-sex-biased genes were characterized by significantly longer distances and lower reliability (lower score and significance) than the linkages to sex-biased genes (Fig. S9).

### Epigenetic features enriched at active sex-biased enhancers

We investigated the enrichment of adult male and adult female mouse liver chromatin states and histone marks (32) in the sets of active, functional sex-biased and cGH-regulated enhancers to identify epigenetic features associated with the regulation of HDI-STARR-seq enhancer activity in a sex-dependent and cGH-responsive manner (Fig. 5). We compared the frequency of each of 14 chromatin states, defined separately for male and female mouse liver (32), across the ATAC regions comprising each sex-biased, cGH-regulated DAR set with that in a background set of 317 robust active but non-regulated ATAC regions (Table S9A, column AJ). Four DAR sets were compared: 41 male-biased and cGH-repressed DARs with regulated activity, and 82 male-biased and cGH-repressed DARs with non-regulated activity; 43 female-biased and cGH-activated DARs with regulated activity, and 53 non-regulated female-biased and cGH-activated DARs (Table S9A, columns U–X). We observed clear differences in chromatin state enrichments between the regulated and non-regulated enhancer DAR sets, with distinct results for the male-biased and the female-biased enhancer DARs. Thus, the 41 regulated male-biased and cGH-repressed DARs were significantly enriched for chromatin state E6, but not for chromatin state E7, in male liver, whereas the 43 regulated female-biased DARs were significantly enriched for chromatin state E7 but not for chromatin state E6 in female liver (Fig. 5A, Table S10A). States E6 and E7 are both characterized by open chromatin and the active enhancer mark H3K27ac; E6 is more highly enriched for the active enhancer mark H3K4me1 while E7 has high probability for the promoter mark H3K4me3 (Fig. 5A; ChromHMM emission probabilities, where DHS indicates open chromatin). Further, in female but not male liver, the set of 41 regulated male-biased and cGH-repressed DARs, but not their non-regulated counterparts (n=82), was significantly enriched for chromatin state E9. State E9 is characterized by the same two active enhancer marks as state E6 – H3K4me1 and H3K27ac – but is devoid of open chromatin. Thus, the male bias and cGH regulation of HDI-STARR-seq enhancer activity of these DARs is significantly associated with the presence of active enhancer marks at the corresponding endogenous mouse genomic sequences in both sexes, but with open chromatin present in male but not female liver. The unique patterns of enrichment of each sex-specifically regulated DAR set for distinct chromatin states in mouse liver support the proposal that the HDI-transfected plasmid DNA sequences acquire histone marks and epigenetic states similar to those found at the corresponding endogenous DNA regions in mouse liver.

**Fig. 5.**
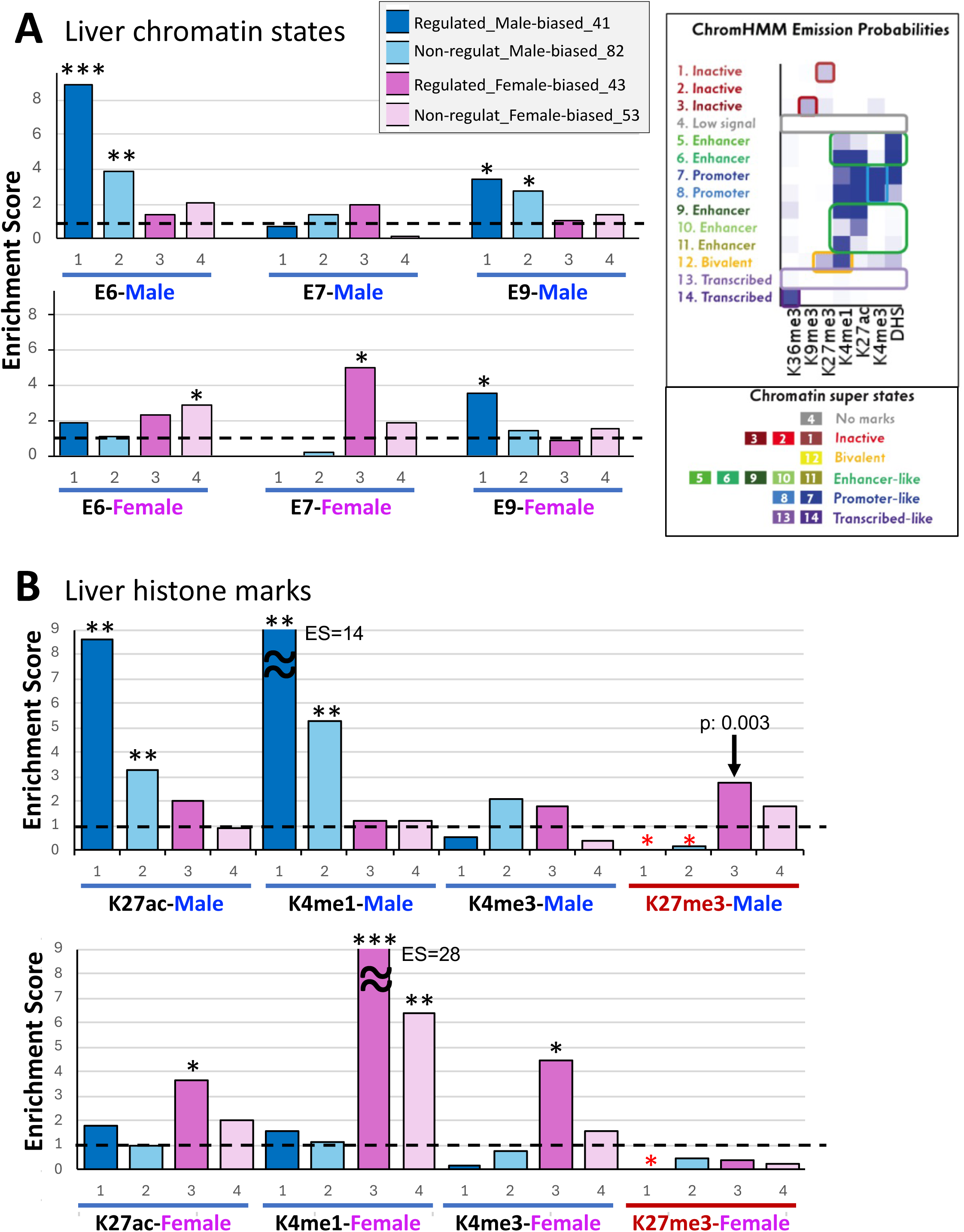
Epigenetic features enriched at regulated versus non-regulated enhancer DARs. Enrichment scores (ES) for the overlap between the indicated sets of sex- and cGH-regulated and non-regulated enhancer DARs identified by HDI-STARR-seq and chromatin states (panel A) and histone marks (panel B) present in mouse liver at the corresponding genomic DNA sequences. Enrichments were calculated for four sets of robust active DAR enhancers: 1) 41 regulated male-biased, cGH-repressed DARs (*dark blue;* also see Fig. 4B, blue box); 2) 82 non-regulated male-biased, cGH-repressed DARs (*light blue*); 3) 43 regulated female-biased, cGH-activated DARs (*dark pink;* also see Fig. 4C, red box); and 4) 53 non-regulated female-biased, cGH-activated DARs (*light pink*). Enrichments were computed relative to a background set of 317 non-regulated genomic regions, which is a subset of 4,126 robust active reporters showing no regulated activity when comparing sexes or the response to cGH-infusion. **(A)** Enrichments for chromatin states E6 and E9 (both are enhancer-like states; see ChromHMM emission probabilities at *right*) and state E7, a promoter-like state, with the chromatin states being those determined in male mouse liver (top row) and those determined in female mouse liver (bottom row). **(B)** Histone-H3 marks identified in male (*top*) or female mouse liver (*bottom*) by ChIP-seq. Horizontal dashed line, ES=1. The significance of enrichment was determined by Fisher’s exact test: *, (−log10P) > 3; **, (−log10P) > 5; ***, (−log10P) > 7, with red asterisks indicating significant depletion. Full datasets, including other chromatin states and other histone marks, are shown in Table S10A.

Consistent with the above findings, the regulated active enhancer DARs were also enriched for genomic regions with specific activating histone marks. The regulated male-biased, cGH-repressed enhancer DARs showed strong enrichments for genomic regions characterized by active enhancer histone marks in male liver (ES = 8.6 for H3K27ac, ES = 14.2 for H3K4me1) but not female liver, with ∼2.7-fold lower enrichments seen for the corresponding non-regulated DARs (Fig. 5B; Table S10A). In contrast, regulated but not non-regulated female-biased, cGH-activated DARs showed strong enrichments for genomic regions characterized by marks for active enhancers (ES = 3.7 for H3K27ac, ES = 27.9 for H3K4me1) and marks for promoters (ES = 4.5 for H3K4me3) in female but not male mouse liver. The regulated male-biased DAR regions were significantly depleted of H3K27me3 repressive histone marks in male liver (ES = 0 to 0.16) (Fig. 5B, *red asterisks*) but showed no significant enrichment or depletion of H3K9me3 or H3K36me3 (repressive and transcribed region marks, respectively; Table S10). In contrast, the set of regulated female-biased, cGH-activated DARs showed weak but significant enrichment for H3K27me3 marks in male liver (ES = 2.75, p = 2.96E-3), which parallels the documented functional repression by H3K27me3 marks of female-biased enhancers in male liver (60).

### Mapping to HDI-STARR-seq active enhancers to disease relevant sex-biased genes

We used multiOmic gene linkage analysis to identify sex-biased genes predicted to be regulated by the 840 DARs targeted by the 1,397 sex-biased and/or cGH-responsive HDI-STARR-seq reporters (Table S9C). 305 of the 840 DARs were linked to genes with corresponding regulation. Thus, 168 male-biased, cGH-repressed DARs were linked to male-biased, cGH-repressed genes, and 137 female-biased, cGH-activated DARs were linked to female-biased, cGH-induced genes (Fig. 4A, *bottom*). Of the other DARs, 37 were sex-biased DARs potentially involved in repressive interactions (31 male-biased, cGH-repressed DARs and 6 female-biased DARs linked to sex-opposite genes), 18 were sex-biased DARs (of which 6 are on ChrX, 6 on ChrY) showing GH-independent sex-biased regulation, and 416 were sex-biased DARs not linked to any genes (n=261) or not linked to any sex-biased genes (n=155) (Table S13D, column K). The male-biased, cGH-repressed DAR gene linkages included MASLD-enabling genes (e.g., *Bcl6*, *Lnc-Lfar1* and *Nox4)*; and the female-biased, cGH-activated DAR linkages included MASLD-protecting genes (e.g., *Hao2*, *Fmo2*, and *Trim24)*. Furthermore, the male-biased, cGH-repressed DARs were primarily in an enhancer state in male liver (n=128 of 168, 76%); the remaining 40 DARs were in promoter (n=18), bivalent (n=6), transcribed (n=12), or inactive (n=4) states. These ATAC regions shifted away from enhancer and promoter states in male liver to increasingly populate bivalent, transcribed and inactive states in female liver (female liver: 102 enhancer (61%), 12 promoter, 11 bivalent, 21 transcribed, 22 inactive) (Table S9C). Similarly, the 137 female-biased DARs showed reciprocal shifts from enhancer and promoter states in female liver to bivalent, transcribed and inactive states in male liver (female: 100 enhancer (73%), 23 promoter (17%), 11 bivalent, 2 transcribed, 1 inactive; male: 71 enhancer (52%), 11 promoter (8%), 24 bivalent, 7 transcribed, 24 inactive) (Table S9C).

### Enrichment of GH-dependent liver TFs at regulated vs. non-regulated sex-biased enhancers

We investigated the regulated sex-biased and cGH-responsive enhancer DARs to determine whether the corresponding genomic sequences show enriched binding by STAT5 and BCL6, two essential, GH-regulated liver TFs implicated in GH-regulated sex-biased liver gene expression (13,32) and in liver metabolic disease (1,5,16,27). We examined the enrichment of sex-biased binding by each TF in regulated versus non-regulated sex-biased DARs compared with a background set of 317 non-regulated enhancers (Fig. 6A). Male-biased STAT5 binding (liver ChIP-seq) (13) was significantly enriched at both regulated and non-regulated male-biased, cGH-repressed DARs. Similarly, female-biased STAT5 binding was enriched at both regulated and non-regulated female-biased, cGH-activated DARs, but was depleted at male-biased, cGH-repressed DARs. Regulated and non-regulated sex-biased, cGH-responsive ATAC sites showed no differences in STAT5 motif frequency (Fig. 6B, Table S13D), suggesting that other features, including histone marks, chromatin states and other bound TFs, are central to the regulation of sex-biased enhancer activity.

**Fig. 6.**
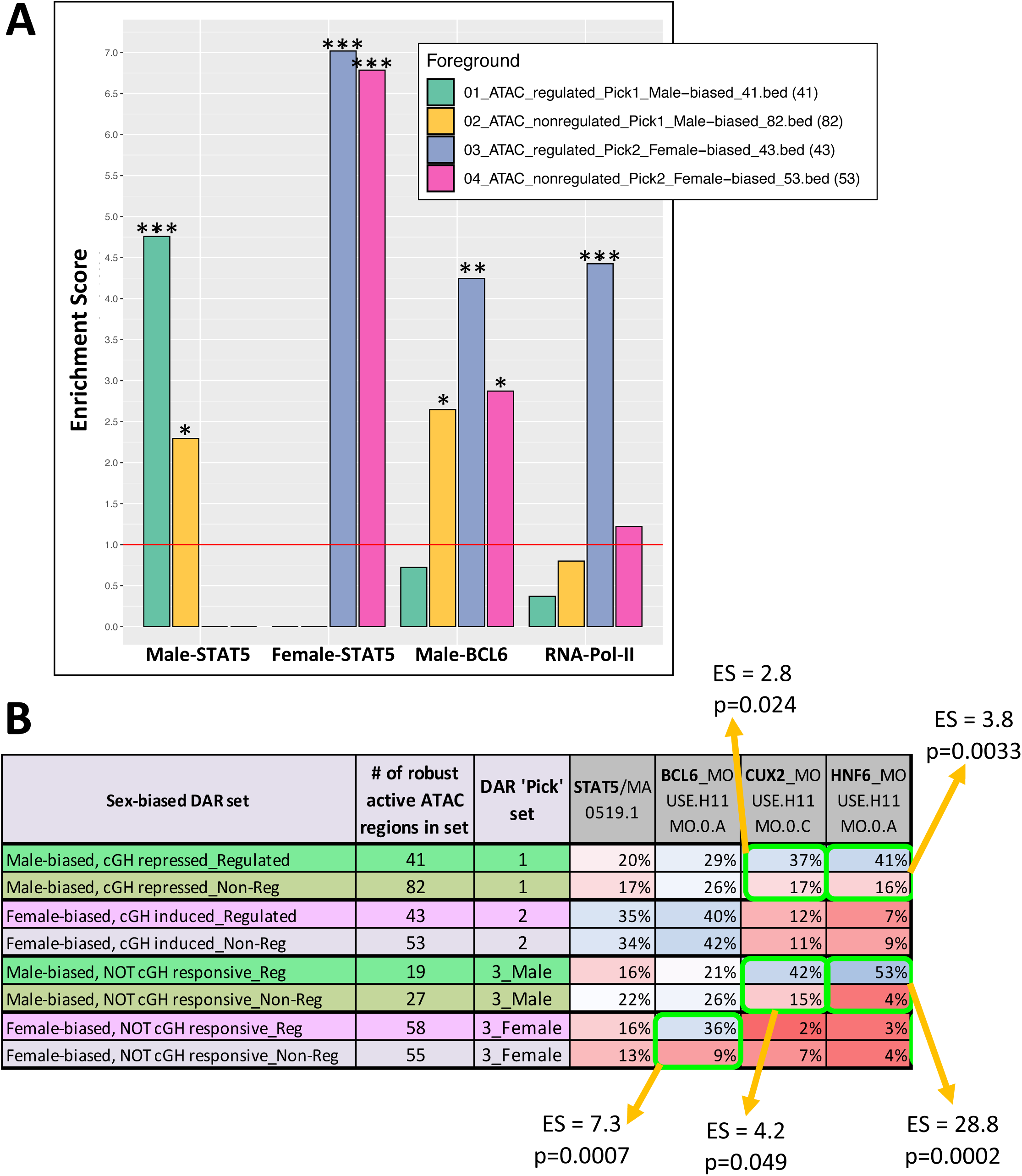
TFs enriched at regulated versus non-regulated enhancer DARs. **(A)** Enrichment scores for the overlap between TF binding sites in mouse liver chromatin and the four indicated sets of sex- and cGH-responsive DARs, grouped by whether the ATAC sites displayed sex- and cGH-regulated enhancer activity by HDI-STARR-seq, or not (‘non-regulated’). STAT5 ChIP-seq sites were either male-biased (n=1,765 sites) or female-biased (n=1,790 sites), as determined by ChIP-seq (13). BCL6 sites: n=6,432 ChIP-seq sites determined in male liver; and RNA-polymerase-II, global liver binding sites (n=31,987). Horizontal red line, ES=1 (no enrichment). The significance of enrichment was determined by Fisher’s exact test: *, p<0.05; **, p<0.01; ***, p<0.001. **(B)** Motif scans for the indicated TF motifs (last 4 columns) list the percentage of each DAR set that contains the motif. Sex-biased DAR sets showing a significant enrichment of the TF motif in the regulated compared to the non-regulated sets are highlighted in green boxes, along with their enrichment scores (ES) and p-values (Fisher’s exact test).

One such factor may be BCL6, whose binding to male mouse liver chromatin was enriched (ES = 4.25, p=1.41E-3) at the set of female-biased, cGH-activated DARs (Fig. 6A), consistent with BCL6 repressing many female-biased genes in male liver (13,32). BCL6 motif frequency was higher in the female-biased than the male-biased DAR sets, but with equal frequency between the regulated and non-regulated sets of sex-biased, cGH-responsive active enhancers (Pick1 and Pick2 sets, Fig. 6B; Table S13D). Of note, BCL6 motifs were significantly enriched at the regulated versus the non-regulated female-biased DARs stringently unresponsive to cGH infusion (Pick3_F DAR set; ES = 7.3, p=0.0007 (Fisher’s Exact test); Fig. 6B). This finding implicates BCL6, a transcriptional repressor showing male-biased expression in liver (26), in controlling sex-biased enhancer activity for these female-biased DARs, whose chromatin accessibility is unresponsive to cGH infusion. The absence of sex-biased enhancer activity at these 55 female-biased DARs can therefore be explained by their deficiency in BCL6 motifs and BCL6-dependent repression in male liver.

Motif scans for CUX2, a female-specific liver TF that represses many male-biased genes in female liver (30), revealed a higher motif frequency in the regulated male-biased versus the regulated female-biased DAR sets (37% vs. 12%, for Pick1 vs. Pick2; 42% vs. 2%, for Pick3_Male vs. Pick3_Female; Fig. 6B). Additionally, CUX2 motifs were significantly enriched in the regulated versus the non-regulated male-biased DAR sets, both those from Pick1 (male-biased DARs suppressed by cGH) and from Pick3_M (male-biased DARs not suppressed by cGH) (ES = 2.8 and 4.2, respectively; Fig. 6B). In view of the near absence of CUX2 protein in male mouse liver (28), these results implicate CUX2 in distinguishing regulated from non-regulated male-biased, cGH-repressed DAR enhancer activity. The motif for HNF6 (Onecut1), which is very similar to that of CUX2 (29), also showed significant enrichment at the regulated versus the non-regulated male-biased DAR sets (Fig. 6B, Table S13D). Together, these findings support the conclusion that the regulated and non-regulated male-biased DARs introduced by HDI bind different subsets of TFs, which are key determinants of the sex bias in enhancer activity detected by HDI-STARR-seq.

### Motif enrichment in sex-biased and cGH-regulated enhancers

We used MEME-CHIP to discover *de novo* motifs for TFs that distinguish regulated from non-regulated sex-biased, cGH-responsive DARs in regulated male-biased, cGH-repressed (n=41) and female-biased, cGH-activated DARs (n=43). The *de novo* motifs identified were matched to a database of established mouse TF motifs (55). Three background sets were used: 1) sex-/cGH-matched ATAC regions with non-regulated enhancer activity; 2) 317 non-regulated robust active ATAC regions; and 3) 65 stringently inactive ATAC regions. Top enriched *de novo* motifs for 41 regulated male-biased, cGH-repressed DARs versus 82 non-regulated male-biased, cGH-repressed DARs were most similar to HNF4A and ESRRA/NR2F2 (Fig. S10). This was confirmed using alternative backgrounds (Table S13B); secondary matches included MAFB and LHX8, in addition to several motifs with no known TF matches. Top enriched motifs for 43 regulated female-biased, cGH-activated DARs versus 53 non-regulated female-biased, cGH-activated DARs best matched SOX7 and CRX_3485.1 (Fig. S11). Secondary matches included SP4/KLF7, NKX2-5, ASCl2 and RFX7 (RFXDC2) (Table S13B).

The *de novo*-discovered motifs were scanned across sex-biased and/or cGH-responsive DARs for frequency differences between regulated and non-regulated enhancer activity DAR sets (Table S13C, Table S13D). Results confirmed significant enrichment of the *de novo* HNF4A-like motif in regulated versus non-regulated male-biased, cGH-repressed DARs (56% vs 15% frequency; ES = 7.45, Fisher’s exact test, p < 0.0001) (Table S13D). Similarly, the *de novo* RFX7-like motif was significantly enriched in regulated versus non-regulated female-biased, cGH-activated DARs (35% vs. 9% frequency; ES = 5.1, Fisher’s exact test, p = 0.0046; Table S13D).

### TAD-based view of active enhancers

We used HDI-STARR-seq to assay 749 ATAC regions across 10 TAD regions (Table S3A, Table S4D). Control TAD regions, devoid of sex-biased genes, generally contained few regulated reporters, as expected. In contrast, the organization of TAD_2885 (chr19; 111 ATAC regions, 47 linked genes, of which 28 are sex-biased) is consistent with domain-level co-localization. This TAD contains the *Cyp2c* gene locus and includes female-biased genes and regulatory regions concentrated at the 5’ end and within a female-biased intra-TAD (35,58), suggesting genuine regulatory activity with tandem reporters showing concordant sex-bias patterns. Additionally, a cGH-suppressed male-biased enhancer near *Cyp2c70* at the 3’ end of TAD_2885, outside the female-biased intra-TAD region, potentially regulates male-biased genes via CTCF/cohesion-mediated loops ∼1 MB to the 5’ end (35). TAD_4231 contains male-biased genes with overlapping correspondingly regulated reporters, confirming sex-bias consistency of the measured HDI-STARR-seq enhancer activity. TAD_4827 is characterized by female-biased genes and female-biased, cGH-activated reporters showing weak STAT5 binding, plus chromatin accessibility differences in ATAC regions showing sex-specific patterns.

### HDI-STARR-seq activity in NPCs

Plasmid DNA introduced by HDI is reported to preferentially transfect hepatocytes versus other liver cell types (61), raising the question of whether NPC enhancer activity can be assayed by HDI-STARR-seq. To address this, we assayed HDI-STARR-seq enhancer activity of 63 ATAC regions (972 reporters) whose chromatin is preferentially accessible in hepatic stellate cells, liver endothelial cells, immune cells, or cholangiocytes (Table S11). Results were compared to a similar number of hepatocyte-enriched ATAC regions. Overall, robust active reporters from ATAC regions with NPC specificity were 1.8-fold depleted (p < 0.00001, Fisher’s exact test) versus hepatocyte ATAC region reporters (Fig. S12A). Depletion was 3.6-fold for hepatic stellate cell-specific reporters (p < 0.0001) and 1.8-fold for liver endothelial cell reporters (p=0.0003) (Table S11). Cholangiocytes, which share many hepatocyte characteristics (36,37), showed a 1.73-fold enrichment of robust active enhancers versus hepatocytes, but this did not reach statistical significance (p = 0.2), probably due to the small number of cholangiocyte-enriched DARs available for analysis. HDI-STARR-seq enhancer activity was significantly higher for hepatocyte ATAC regions than for NPC ATAC regions (Fig. S12C), consistent with the hepatocyte preference of HDI.

## Discussion

This study establishes a functional genomics framework that moves beyond correlative chromatin maps to identify and then directly validate in intact mouse liver hundreds of sex-biased, GH-responsive hepatocyte enhancers driving liver sexual dimorphic gene expression, including expression of genes associated with sex differential, GH-regulated MASLD susceptibility. By integrating hepatocyte-resolved single-nucleus multiOmics for both male and female mice (40) with functional enhancer profiling in intact liver using HDI-STARR-seq (41), we experimentally validated sex-biased, GH-regulated reporters representing 840 enhancer DARs, including 168 male-biased and cGH-repressed DARs and 137 female-biased and cGH-activated DARs. Critically, our *in vivo* approach overcomes critical limitations of prior studies by assaying enhancer function in intact liver under physiological conditions, where transfected episomal reporters undergo chromatinization (41,62,63) and respond to endogenous regulatory factors within their native hormonal and metabolic environment. Together, our findings establish three major mechanistic principles, discussed below, that advance our understanding of how GH-regulated chromatin accessibility drives sex-dependent hepatocyte gene expression linked to metabolic disease risk.

First, sex-biased hepatocyte chromatin accessibility is a strong but imperfect predictor of functional sex-biased enhancer activity. While 63% of DNA-qualified reporters (11,996 of 19,136) showed enhancer activity under at least one biological condition, a smaller fraction exhibited regulated enhancer activity in response to sex or GH signaling. This is consistent with our tiling design, in which each ATAC region was covered by multiple overlapping 180-nt reporters spaced 55 bp apart. Functional enhancer activity is often restricted to discrete sequence elements within accessible chromatin regions, and thus only a subset of reporters tiling a regulated DAR would be expected to show regulated activity. Indeed, Many DARs gave concordant regulatory patterns across neighboring reporters, supporting the presence of localized functional elements embedded within broader accessible chromatin regions. Moreover, analysis at the ATAC region (DAR) level revealed a highly significant correlation between sex-bias and cGH-responsiveness of enhancer activity (Fig. 4D). Furthermore, we identified 41 male-biased, cGH-repressed DARs harboring regulated reporters showing matched male-biased, cGH-repressed enhancer activity, and similarly for 43 female-biased, cGH-activated DARs with matched female-biased, cGH-induced enhancer activity, demonstrating that concordance between chromatin state and regulation of enhancer activity is non-random. However, these matching patterns were observed for only a subset of the sex-biased and cGH-regulated hepatocyte DARs, indicating that open, accessible chromatin is a necessary but not sufficient condition for enhancer function. This finding reconciles earlier genome-wide DNase-seq studies, which identified thousands of sex-biased DHS sites in bulk liver tissue that co-localize with or are found nearby sex-biased genes (31,32), with the reality that only a subset of accessible regions function as bona fide enhancers *in vivo* under physiological conditions. The discordance between chromatin accessibility and enhancer function underscores the importance of direct functional validation and indicates that additional determinants, such as specific TF binding motifs, cooperative TF occupancy, histone modifications, and three-dimensional chromatin architecture, are required to convert an accessible chromatin region into an active enhancer, in accord with results from massively parallel reporter assays in other biological systems (64–66). Importantly, our tiling approach, while enabling fine mapping of functional sequences within DARs, may lead to false negatives when critical enhancer elements span reporter boundaries or require longer sequence contexts for full activity. Nevertheless, our study extends these general principles to sex-biased hepatocyte enhancers assayed in intact mouse liver under conditions where episomal reporters become chromatinized and respond to endogenous hormonal cues.

Second, functional sex-biased, GH-responsive enhancers are distinguished from non-functional sex-biased DARs by distinct epigenetic signatures that predict responsiveness to hormonal regulation. Among male-biased, cGH-repressed DARs, those showing correspondingly regulated HDI-STARR-seq enhancer activity were highly enriched for genomic sequences characterized by chromatin state E6 with its dual enhancer histone marks in male mouse liver (H3K4me1, 14.2-fold; H3K27ac, 8.6-fold) compared to a background set of 317 robust active but non-regulated genomic regions. Similarly, female-biased, cGH-activated DARs showing correspondingly regulated enhancer activity were strongly enriched for genomic sequences with enhancer marks in female mouse liver (H3K4me1, 27.9-fold; H3K27ac, 3.7-fold) but in addition showed a unique 4.5-fold enrichment for both the promoter mark H3K4me3 (chromatin state E7) and RNA polymerase-II-bound genomic regions, suggesting that these enhancers — which are not closer to their linked gene targets than male-biased enhancers — adopt promoter-like regulatory architectures. Strikingly, regulated male-biased enhancer DARs were depleted of H3K27me3 repressive histone marks in male liver, while regulated female-biased enhancer DARs were enriched for H3K27me3 marks in male liver, consistent with the H3K27me3-mediated functional repression of female-biased genes established in male liver (60). Taken together, these findings provide strong support for the proposal that episomal HDI-STARR-seq reporters delivered to mouse liver acquire endogenous hepatocyte-like epigenetic states in a sequence-dependent manner, enabling functional enhancer activity when the appropriate activating marks are present and repressive marks are absent. Further, our findings indicate that enhancer sequences 180 bp in length (Fig. 2C), when combined with the minimal *Alb* promoter incorporated in our HDI-STARR-seq reporter plasmids, encode sufficient information to recruit hepatocyte chromatin modifiers, and perhaps could be employed to study the impact on sex-biased enhancer function and sex-differential disease susceptibility when these epigenetic states are disrupted by genetic variants (34) or environmental exposures (67).

Third, binding site architecture for GH-regulated TFs discriminates functional, regulated enhancers from non-functional DARs with similar chromatin accessibility. Sex-biased STAT5 binding to liver chromatin was significantly enriched at the correspondingly sex-biased and cGH-responsive DAR sequences but with no large differences between regulated and non-regulated DAR enhancers. This may, in part, reflect the direct role that STAT5 plays in sex-biased chromatin opening *per se* for a subset of male-biased open chromatin regions (33). In contrast, motifs for the female-specific transcriptional repressor CUX2 (30) were enriched at regulated compared to non-regulated male-biased DARs, while motifs for the male-biased transcriptional repressor BCL6 (13,26,27) were enriched at regulated versus non-regulated female-biased DARs unresponsive to cGH. These patterns suggest a model in which STAT5 binding at male-biased DARs is necessary but not sufficient for regulated enhancer activity. Hormonal responsiveness of DAR enhancer activity would be conferred by additional determinants, such as CUX2 motifs that enable female-specific repression regulated by cGH, and BCL6 motifs that enable male-specific repression of female-biased enhancers. *De novo* motif discovery supported this model, identifying HNF4A-like motifs enriched at regulated versus non-regulated male-biased DARs and RFX7-like motifs enriched at regulated female-biased DARs. HNF4A is a liver-enriched transcription factor that cooperates with STAT5 to activate male-biased genes (25) and its motif enrichment at functional enhancers suggests it acts as a co-determinant of GH-responsive enhancer activity. Motif enrichment for RFX7 at regulated female-biased enhancers nominates this tumor suppressor (68) as a novel candidate for female-biased hepatic gene regulation. These findings further refine the current model of GH-regulated liver sexual dimorphism: rather than STAT5 and STAT5-regulated repressors (BCL6, CUX2) acting in simple opposition, our data suggest a layered regulatory code in which combinatorial motif architecture—STAT5 plus cooperating activators (HNF4A) and/or sex-biased repressors (BCL6, CUX2)—determines which accessible chromatin regions convert GH input signals into functional, hormone-responsive enhancer output.

Integration of functional enhancer datasets with sex-biased genes revealed mechanistic links between sex-biased, GH-regulated enhancers and sex-differential MASLD susceptibility. Of 840 functional enhancer DARs showing sex-biased or cGH-responsive enhancer activity, 168 were male-biased, cGH-repressed DARs linked to correspondingly regulated male-biased genes and 137 were female-biased, cGH-activated DARs linked to correspondingly regulated female-biased genes. Critically, a subset of the male-biased enhancer DARs linked to known MASLD-enabling genes, including *Bcl6* (male-biased transcriptional repressor that enhances MASH progression by repressing PPARA-dependent hepatoprotective fatty acid oxidation genes (5,27,69)), *Nox4* (NADPH oxidase that prevents oxidative damage in early stages of disease (70) but become maladaptive and mediates oxidative stress, inflammation, and fibrosis as disease advances (71)), and the lncRNA lipid metabolism regulator *Lnc-Lfar1* (72). Similarly, female-biased enhancer DARs were linked to MASLD-protective genes, notably *Hao2* (hydroxyacid oxidase promoting fatty acid oxidation linked to tumor suppression (73,74)), *Fmo2* (flavin monooxygenase with anti-inflammatory roles that inhibits SREBP1 translocation, blocking steatosis (75)) and *Trim24* (epigenetic regulator that represses steatosis, inflammation and fibrosis (76,77)). Notably, chromatin state analysis showed that the male-biased enhancers shift from active enhancer and promoter states in male liver to increasingly populate inactive, bivalent, or transcribed states in female liver, consistent with epigenetic silencing of male-biased disease-promoting pathways in females. Conversely, female-biased enhancers tend to shift from active states in female liver to more repressed states in male liver, consistent with loss of protective epigenetic modifications in males. These findings provide a mechanistic framework for understanding the higher MASLD prevalence in males and post-menopausal females (1) and suggest therapeutic strategies targeting GH signaling or enhancer epigenetic states to modulate sex-biased metabolic disease risk. Moreover, identification of functional enhancers linked to *Bcl6,* a male-biased, cGH-repressed regulator balancing immunity and metabolism and whose disruption drives fatty liver specifically in males (5,27,69), suggests that targeting BCL6-regulated enhancer networks may offer sex-specific therapeutic opportunities.

The HDI-STARR-seq platform circumvents limitations of cell culture-based massively parallel reporter assays (MPRAs) and enables massively parallel functional assays of enhancer activity within hepatocytes *in situ*, where episomal reporters apparently undergo chromatinization, recruit endogenous TFs and, as we establish here, respond to plasma GH pattern-dependent hormonal signaling. Unlike traditional transgenic enhancer-reporter assays, which are limited to single enhancers and require extensive mouse breeding, HDI-STARR-seq enabled the parallel testing of 1,839 ATAC regions (23,912 tiled reporters) in individual mice, dramatically accelerating enhancer discovery and characterization. The 11-day interval between HDI and liver tissue collection ensured recovery from the initial stress associated with HDI (61,78), as verified by absence of stress gene induction, while retaining robust plasmid expression and sufficient time for cGH-driven chromatin remodeling and epigenetic reprogramming. Importantly, our finding that the robust active set of HDI-transfected regulated enhancers shows strong, significant enrichment for genomic regions whose endogenous liver loci have active histone marks and are in active enhancer chromatin states with depletion of repressive histone marks provides strong support for the proposal that the transfected episomal DNA retains sufficient *cis*-regulatory sequence information to recruit endogenous TFs and their associated transcriptional regulatory complexes and chromatin modifiers to recreate native-like chromatin environments. Of note, a potential limitation of HDI-STARR-seq is the preferential transfection of hepatocytes by HDI (78), which is supported by our finding of an up to 3.6-fold depletion of robust active enhancers in NPC–specific ATAC regions. However, this likely has limited impact on our conclusions, given that the large majority of sex-biased, cGH-regulated DARs and genes are almost exclusively limited to hepatocyte clusters (4,40). Future adaptations of HDI-STARR-seq using cell-type–targeted viral delivery methods or lipid nanoparticles may extend this method to NPCs.

Our findings also have important implications for understanding hormone-regulated gene expression and for developing sex-specific therapeutic strategies. The principle that chromatin accessibility is necessary but insufficient for enhancer function raises the possibility that genome-wide association studies identifying disease-associated SNPs in accessible chromatin may substantially overestimate the fraction of variants that directly alter enhancer function. Functional validation using HDI-STARR-seq or similar in vivo assays will be essential to prioritize variants for mechanistic follow-up. Further, our discovery that specific TF motifs, such as HNF4A, discriminate regulated from non-regulated enhancers identifies these factors as promising candidates for future mechanistic investigation. Moreover, our finding that regulated female-biased enhancer DARs are specifically enriched for repressive H3K27me3 marks in male liver suggests a mechanism for epigenetic suppression of female-typical enhancer activity in males and the possibility that interventions blocking H3K27me3 deposition or activating H3K27me3 demethylases at these enhancers in males could confer metabolic protection. Finally, extending HDI-STARR-seq to widely studied liver disease models, including high-fat diet, methionine-choline–deficient diet and genetic obesity models (79), may identify enhancers whose activity is dysregulated by metabolic stress, revealing how environmental and genetic risk factors interact with sex-biased regulatory programs.

### Study limitations

We recognize several limitations. First, our 180-nt reporter design with 55 bp overlapping reporters, while enabling fine mapping within ATAC regions, may fragment longer enhancers and miss cooperative interactions between distal elements. Future studies may investigate the extent to which hepatocyte enhancer activity is enhanced by extending sequence context using full-length ATAC regions (mean length, 634 bp) as single reporters. Second, by coupling all reporters to the liver-specific *Albumin* minimal promoter (41) we may bias detection toward enhancers compatible with this promoter architecture; testing reporters with alternative minimal liver promoters could reveal promoter-specific enhancer–promoter compatibility issues or biases. Third, the multiOme peak-to-gene linkage analysis we employed predicts enhancer–gene relationships based on correlations, which do not prove causality; definitive validation will require CRISPR-mediated deletion of individual enhancers *in vivo* and measurement of target gene expression, e.g., using AAV-delivered Cas9 and multiplexed gRNA libraries. Fourth, as HDI-STARR-seq interrogates enhancer activity on episomal plasmids, our data do not establish necessity or sufficiency at native chromosomal loci. Finally, our finding that 37% of DNA-qualified STARR-seq reporters showed no enhancer activity in any of three biological conditions suggests either that many accessible chromatin regions are not enhancers, or that they require specific TFs, co-factors, metabolic states or promoter architectures not represented under our experimental conditions; testing reporters under diverse physiological perturbations (fasting, high-fat diet, xenobiotic exposures) may activate currently silent enhancers and reveal context-dependent regulatory logic.

In summary, this study establishes HDI-STARR-seq as a powerful *in vivo* platform for systematically validating hormone-regulated enhancers and defines a functionally validated enhancer atlas linking GH-regulated chromatin accessibility to hepatocyte expression of many sex-biased genes, including genes impacting MASLD susceptibility. Our findings that only a minority of sex-biased DARs function as regulated enhancers, that functional enhancers are distinguished by combinatorial epigenetic and TF codes, and that these enhancers are linked to disease-relevant metabolic pathways, provide a mechanistic framework for understanding how hormonal cues—encoded in pulsatile versus continuous pituitary GH secretion patterns—are decoded by the genome into sex-specific transcriptional programs and sex-differential disease outcomes. Together, these insights establish a foundation for future studies aimed at elucidating genetic and environmental perturbations that disrupt sex-biased enhancers and uncover novel therapeutic targets within enhancer-regulated pathways contributing to sex differences in liver metabolism and disease susceptibility.

## Acknowledgments

The authors thank Drs. Maxim Pyatkov and Kritika Karri for advice and collaborative assistance and Drs. Christine Goldfarb and Ravi Sonkar for assistance with tail vein injections.

## Data availability

All Raw fastq sequencing files are available at GEO (https://www.ncbi.nlm.nih.gov/geo/) under accession number GSE315771. Other data generated and analyzed during this study are included in this published article or in the data repositories listed in References.

## Author contributions

This study was jointly designed by TYC and DJW. TYC carried out all wet lab work, including HDI-STARR-seq library preparation, HDI-STARR-seq mouse studies, and Illumina sequencing library preparation, validation and initial quality control analysis. TYC carried out primary analysis of the HDI-SATARR-seq data, and TYC and DJW jointly performed secondary data analysis, figure preparation and preparation of Supplementary tables and other Supplementary files for publication. TYC wrote a first draft of the manuscript, which was revised, edited and finalized by DJW. DJW supervised the overall project.

## Supplemental Figure legends

**Fig. S1.**
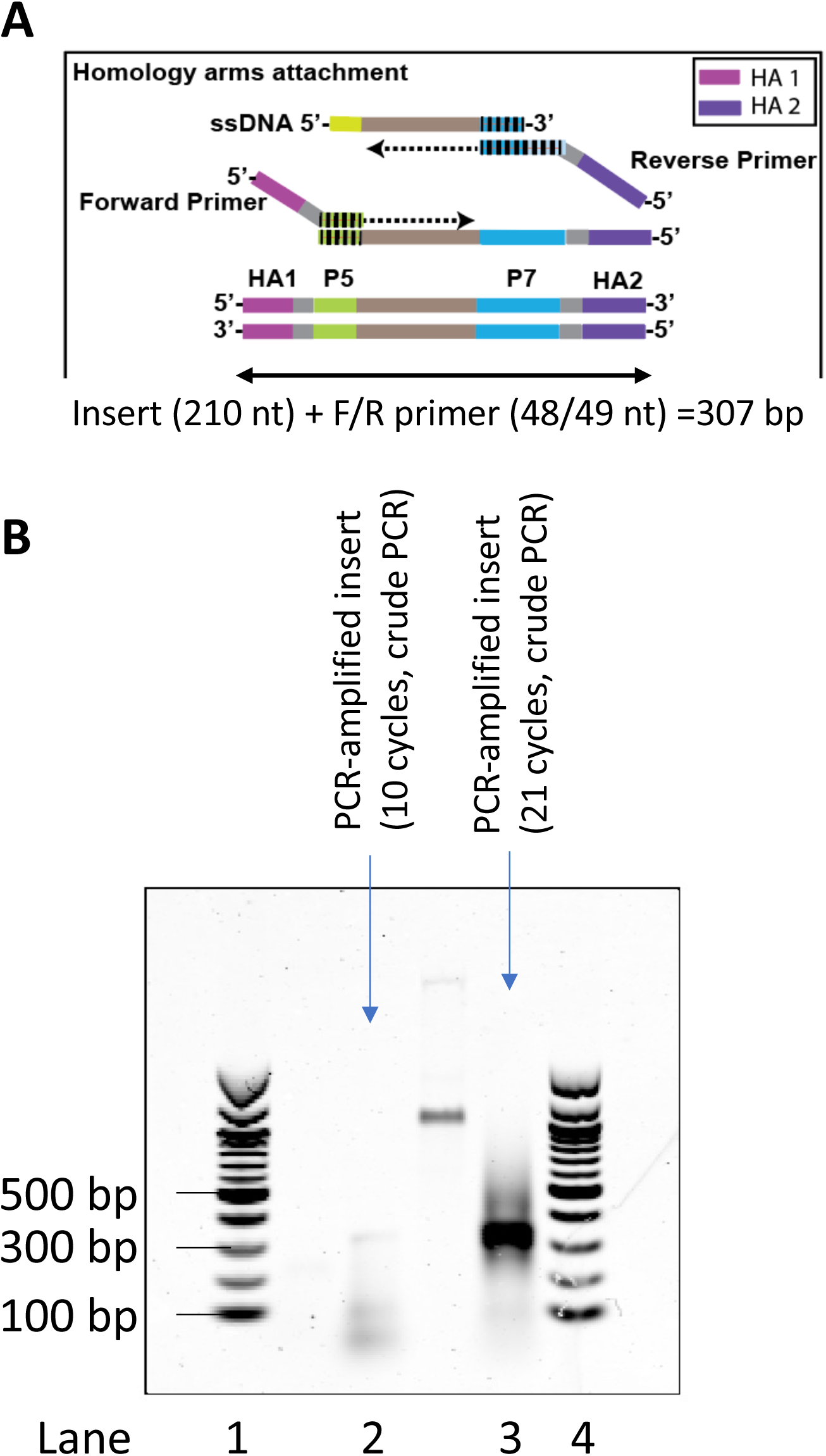
Synthetic STARR-seq preparation. (A) Pooled inserts, started from synthetic ssDNA oligos, were amplified by PCR to attach sequences of homology arms at ends, were expected to be 307 bp long. (B) Crude PCR products with 10 and 21 PCR cycles. Gel electrophoresis showed the size of the PCR fragments after 10 amplification cycles (Lane 3) and 21 amplification cycles (Lane 5) matched with the expected length. The fragments with 10 PCR cycles were proceeded for plasmid library preparation. Lanes 1 and 4 are 100 bp NEB marker (New England Biolabs, #N3231S). Lane 2 represents the amplified HA-adapted DNA fragments when minimizing the numbers of PCR cycles, conditions that are used in library preparation. The lower molecular bands indicate unused primers and primer dimer, to be removed using SPRI beads prior to the subsequential cloning steps. Lane 3 shows the pattern observed when STARR-seq inserts are over-amplified.

**Fig. S2.**
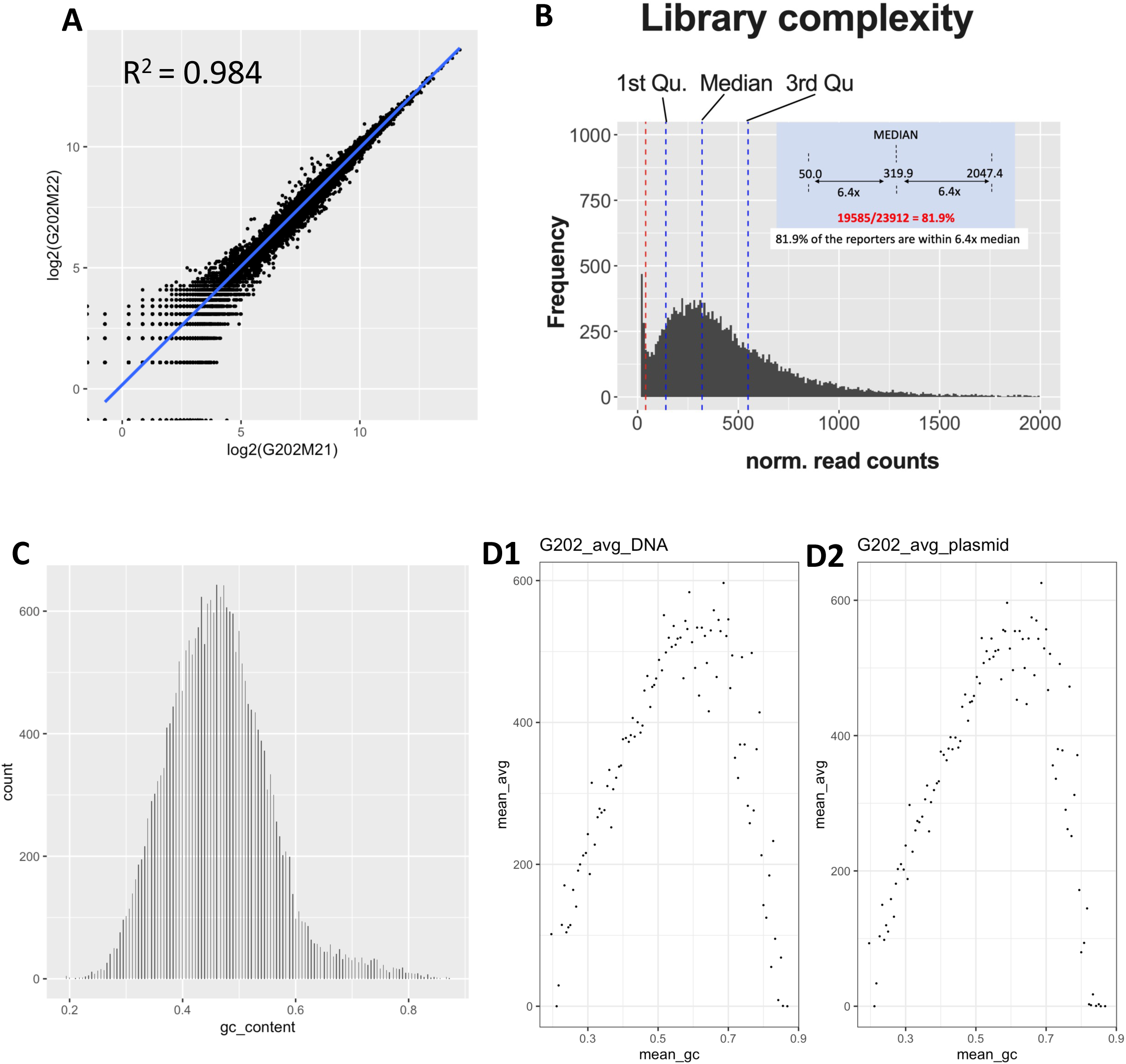
Reporters cloned into HDI-STARR-seq library: characterization with respect to GC content. (A) Scatterplot showing alignment of normalized read counts in two technical replicates of the plasmid libraries amplified separately (Illumina sequencing libraries G202M1 and G202M2; see Table S6). (B) Frequency distribution of averaged normalized read counts within 23,912 reporters from 2 plasmid libraries and 3 DNA libraries. 81.9% of reporters are within the range of 6.4x median. (C) Frequency distribution showing the GC content of 23,912 individual synthetic reporters (based on the 180 nt sequence). Y-axis, number of library reporters at each % GC that were included in the design for the set of 23,912 reporters. (D) Averaged read counts of 23,912 reporters from 5 plasmid/DNA libraries (including 2 plasmid (D1) and 3 extracted DNA (D2) libraries) were sorted by GC content, then grouped into 500 bins. Data show biphasic correlation of reporter representation as judged by read counts in PCR-amplified libraries versus GC content.

**Fig. S3.**
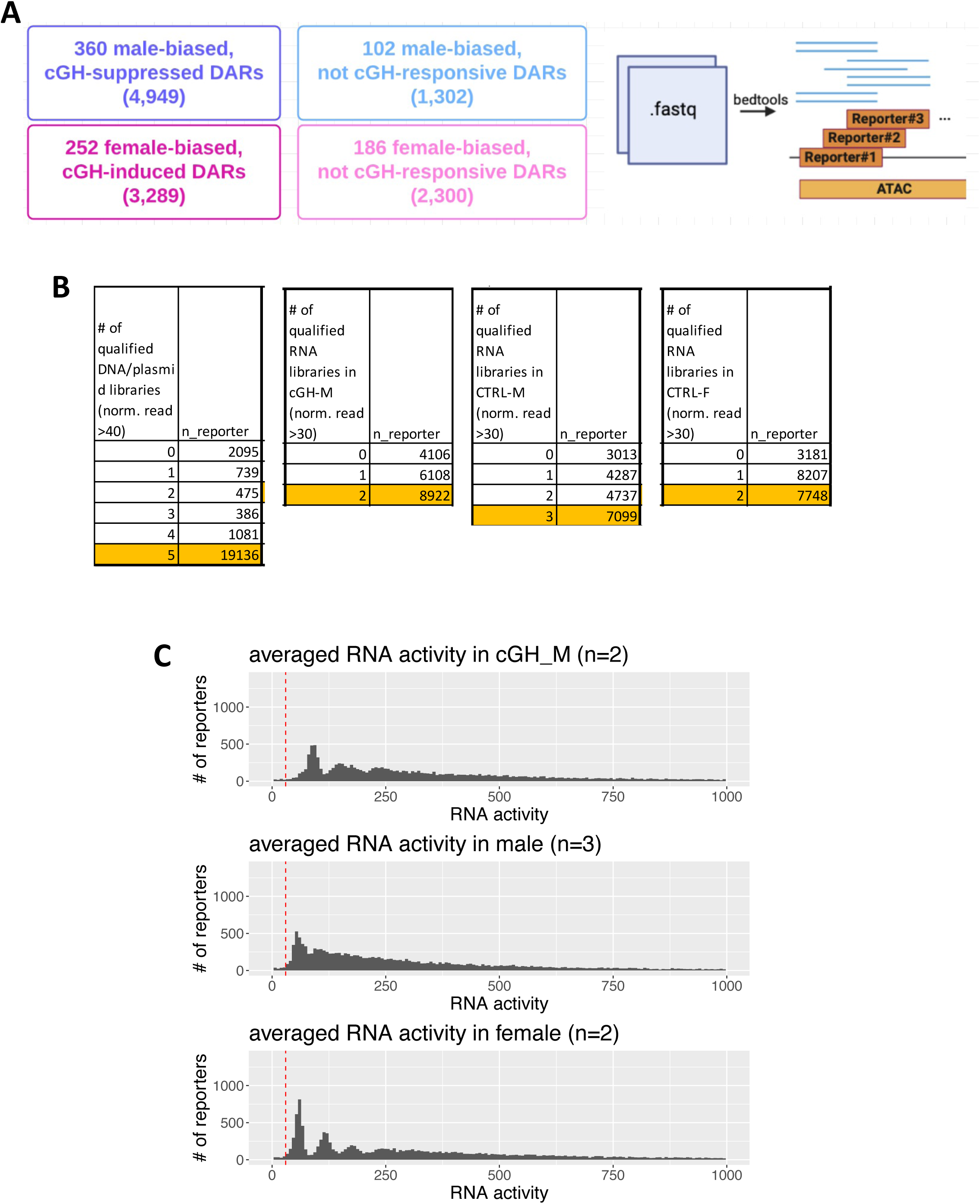
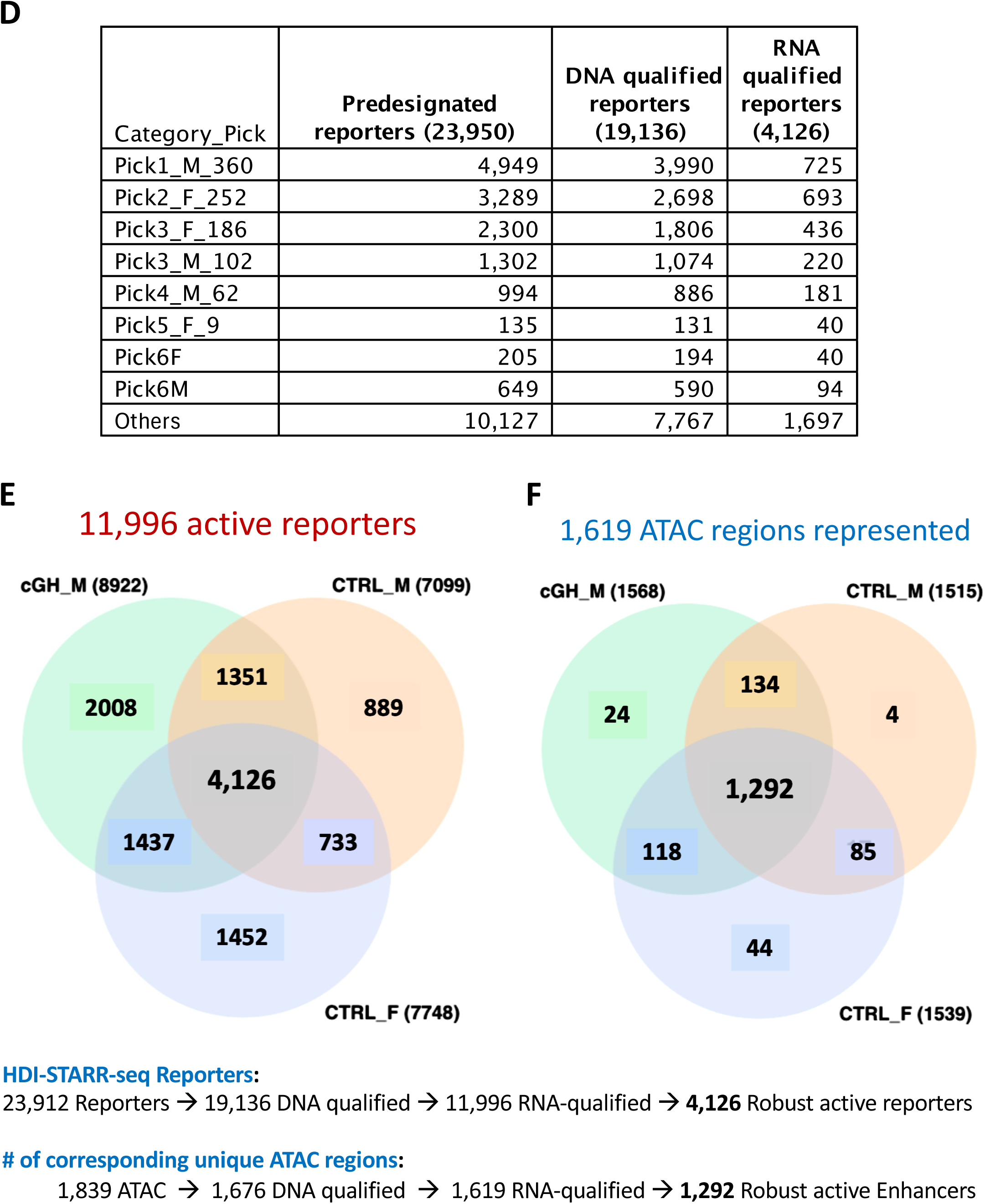
Active reporters identified in mouse liver by HDI-STARR-seq across three biological conditions. (A) 900 selected sex-biased DARs were tiled with reporters (numbers in parenthesis, e.g., 4,949 reporters designed for 360 male-specific, cGH-repressed DARs). Read count for each reporter was calculated based on >95% overlap with the genomic coordinates of 23,912 reporters. (B) The first table shows the number of DNA or plasmid libraries (out of the 5 such libraries examined) that met the threshold of >= 40 normalized reads per 10 million mapped reads for each DNA or plasmid library. Thus, 19,136 reporters were so qualified in all 5 plasmid + DNA libraries, 1,081 other reporters were qualified in only 4 of the 5 libraries, etc. The 3 other tables indicate the number of active reporters (out of the 19,136 reporters qualified based ion DNA) in each of the three biological conditions based on the normalized read counts in either both (n=2) or all 3 (n=3) of the liver RNA libraries analyzed by HDI-STARR-seq for that biological condition. For the RNA libraries, ‘qualified’ indicates the liver met the threshold of 30 normalized reads per 10 million mapped reads for STARR-seq RNA libraries recovered from the HDI-transfected livers. (C) Histograms showing average RNA activity of 19,136 DNA qualified reporter regions in cGH-M (n=2), male (n=3) and female (n=2) mice, grouped into 150 bins. (D) Table showing the total number of reporters designed, the number of reporters within the HDI-STARR-seq library that were qualified (DNA qualified) and the numbers of reporters that were RNA qualified after HDI-STARR-seq for each of the indicated DAR categories selected for HDI-STARR-seq analysis (Table S2). (E, F) Venn diagrams showing the 11,996 active reporters from the union of three active sets in three biological conditions (8,922 active reporters in cGH-treated male livers, 7,099 active reporters in male livers and 7,748 active reporters in female livers), and the number of ATAC regions represented by each subset. 4,126 reporters are robustly active in common in all three biological conditions in terms of RNA qualification and DNA qualification and comprise 1,292 unique ATAC regions. See Table S7A. <u>Note</u>: Although, for example, there are 2,008 active reporters that are uniquely active in cGH-treated male liver (panel E), these reporters map to a total of 1,063 of the 1,292 ATAC regions represented by at least 1 of the 4,126 robust active reporters. Thus, many of these 1,063 ATAC regions have one or more other individual reports that are robust active reporters. Consequently, there are only n=24 ATAC regions that harbor reporters active in only cGH-infused mouse liver.

**Fig. S4.**
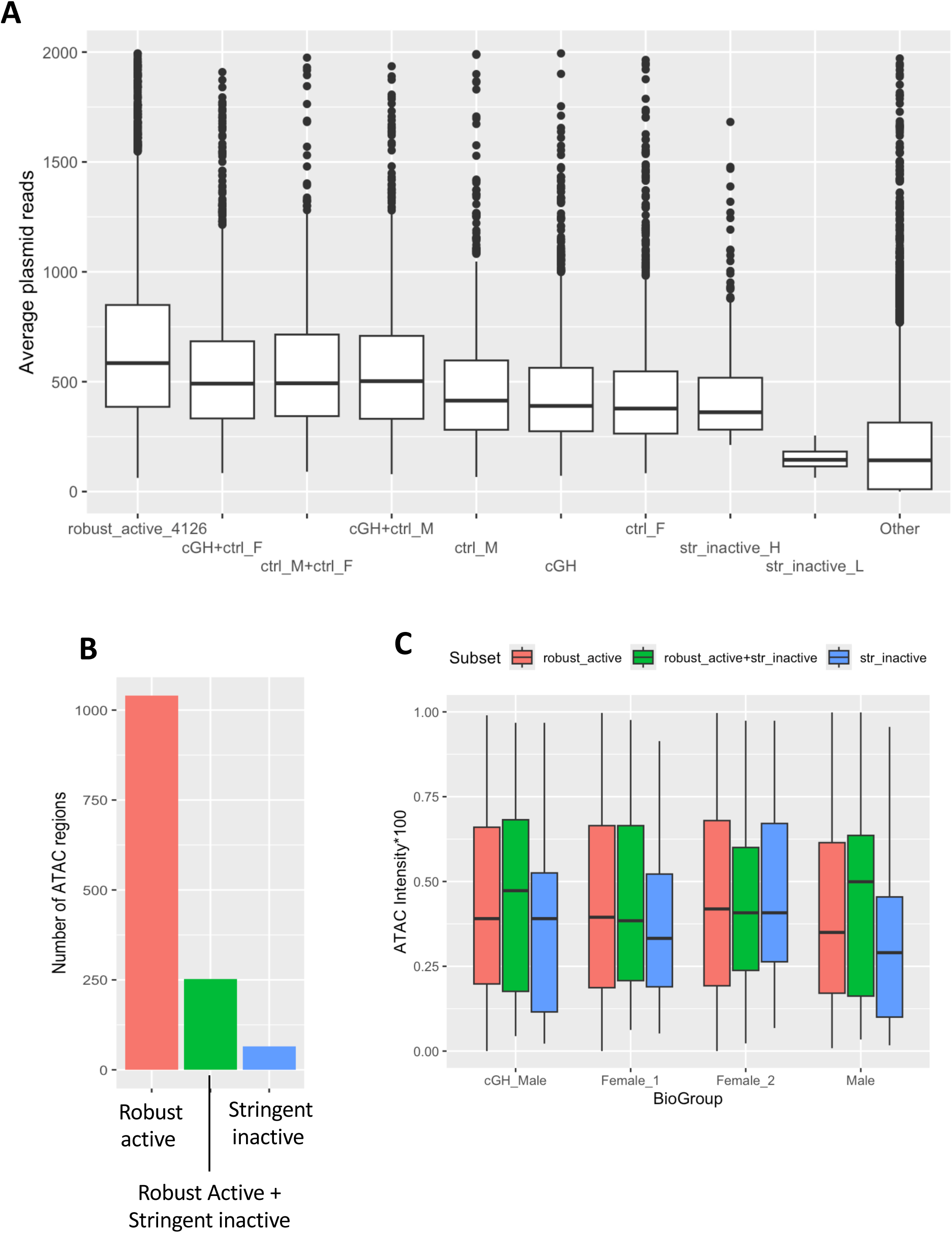
Active reporters and stringently inactive reporters: plasmid library representation, number of ATAC regions, and ATAC intensity values. (A) Plasmid reads in sets of 4,126 robust active reporter regions whose recovered liver RNA met the threshold for qualification in both replicate livers (cGH-treated male, female livers) or in all 3 replicate livers (male) have the highest representation in the original STARR-seq plasmid library used for HDI. The set of 982 stringently inactive reporters are defined as the regions consistently exhibiting low activity (less than 30 normalized reads) in RNA libraries of all biological replicates. We defined the top 400 stringently inactive reporters with higher plasmid representation (labeled as “str_inactive_H”) and the other 582 stringently inactive reporters with lower plasmid reads (labeled as “str_inactive_L”). The set of 582 stringently inactive reporters was excluded from downstream comparisons to avoid artificial effect of the low representation in the library. (B) Number of ATAC regions represented by robust active reporters only, by a mixed of robust active and by stringently inactive reporters only. (C) Boxplots show the ATAC cut sites intensity distributions (ATAC intensity*100) in hepatocytes for the three subsets in panel B from the four biological samples analyzed by MultiOmics. Result shows that stringent inactive set_H is comparable as a background set for enrichment analyses, in terms of its plasmid representation and chromatin accessibility in hepatocytes.

**Fig. S5.**
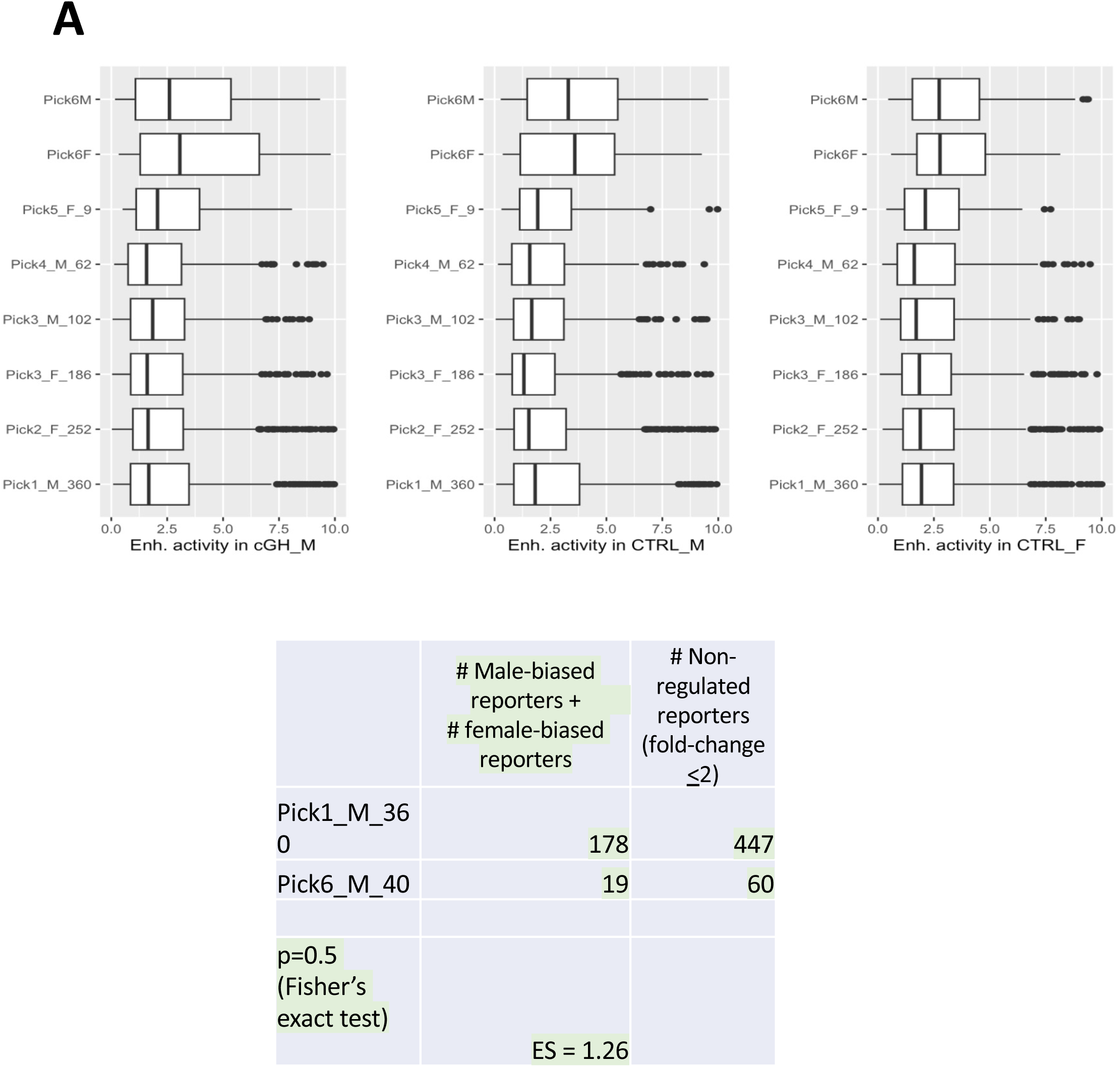
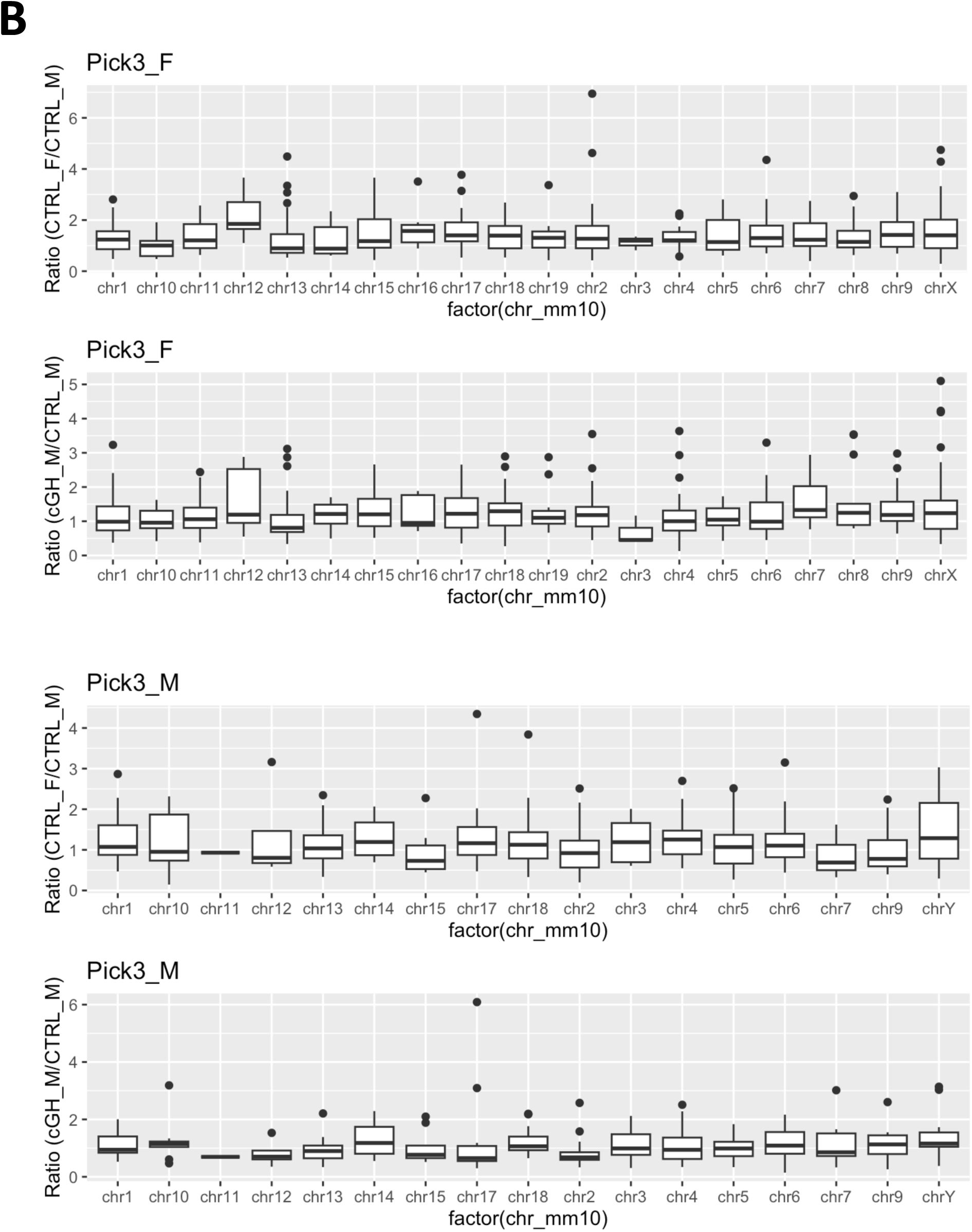
HDI-STARR-seq reporter activity for ATAC regions that are far from genes (A) and are located on autosomal vs sex-chromosomes (B). (A): Boxplots (top) show the distribution of STARR-seq activity for all three biological conditions for individual robust active reporters in each of the 6 indicated DAR Pick categories. Results indicate that DAR regions in Pick6, which are distant from genes, are active, and the table at the bottom shows that Pick1 and Pick6 are very similar regarding their frequencies of robust active reporters. Based on enrichment analysis using Fisher’s exact test, there is no enrichment of robust active reporters in Pick6 as compared to robust active reporters in Pick1. (B): Boxplots showing the distributions of enhancer activity (female/male liver ratio, and cGH-infused male/male liver ratio, as indicated) for each mouse chromosomes for all robust active reporters from Pick3_F and Pick3_M. The data shown represent 436 and 220 robust active STARR-seq probes in Pick3_F and Pick3_M, respectively. No difference in STARR-seq reporter activity was seen for sex chromosomes as compared to autosomal chromosomes, by judged from Kolmogorov-Smirnov test. Almost all STARR-seq reporters assaying sex chromosome are in Pick3_F and Pick3_M, except 1 reporter from Pick6.

**Fig. S6.**
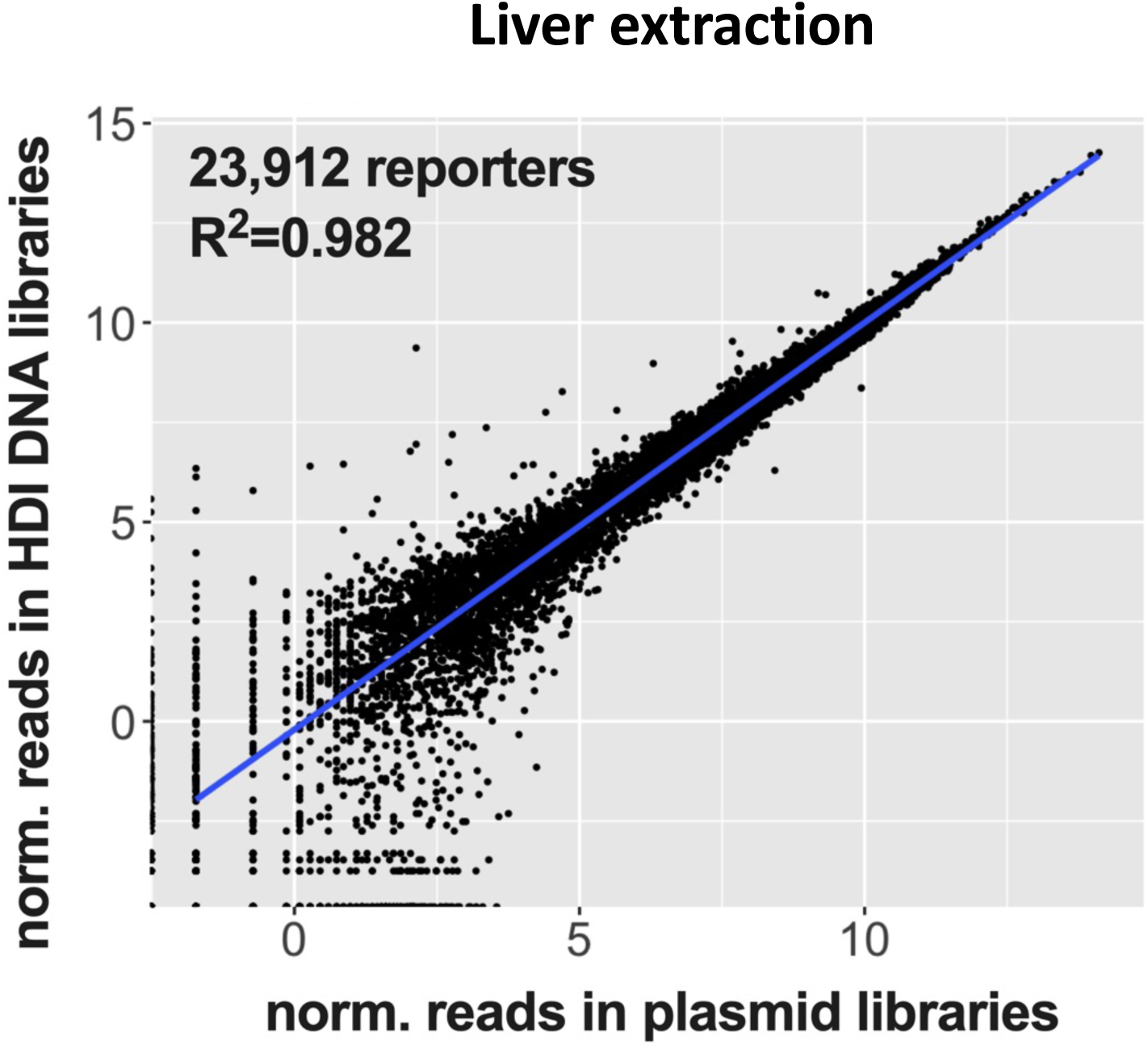
Liver-extracted HDI-STARR-seq DNA library reads correlate well with HDI-STARR-seq plasmid library reads, indicating no significant reporter sequence-dependent bias in plasmid uptake or retention by hepatocytes following HDI.

**Fig. S7.**
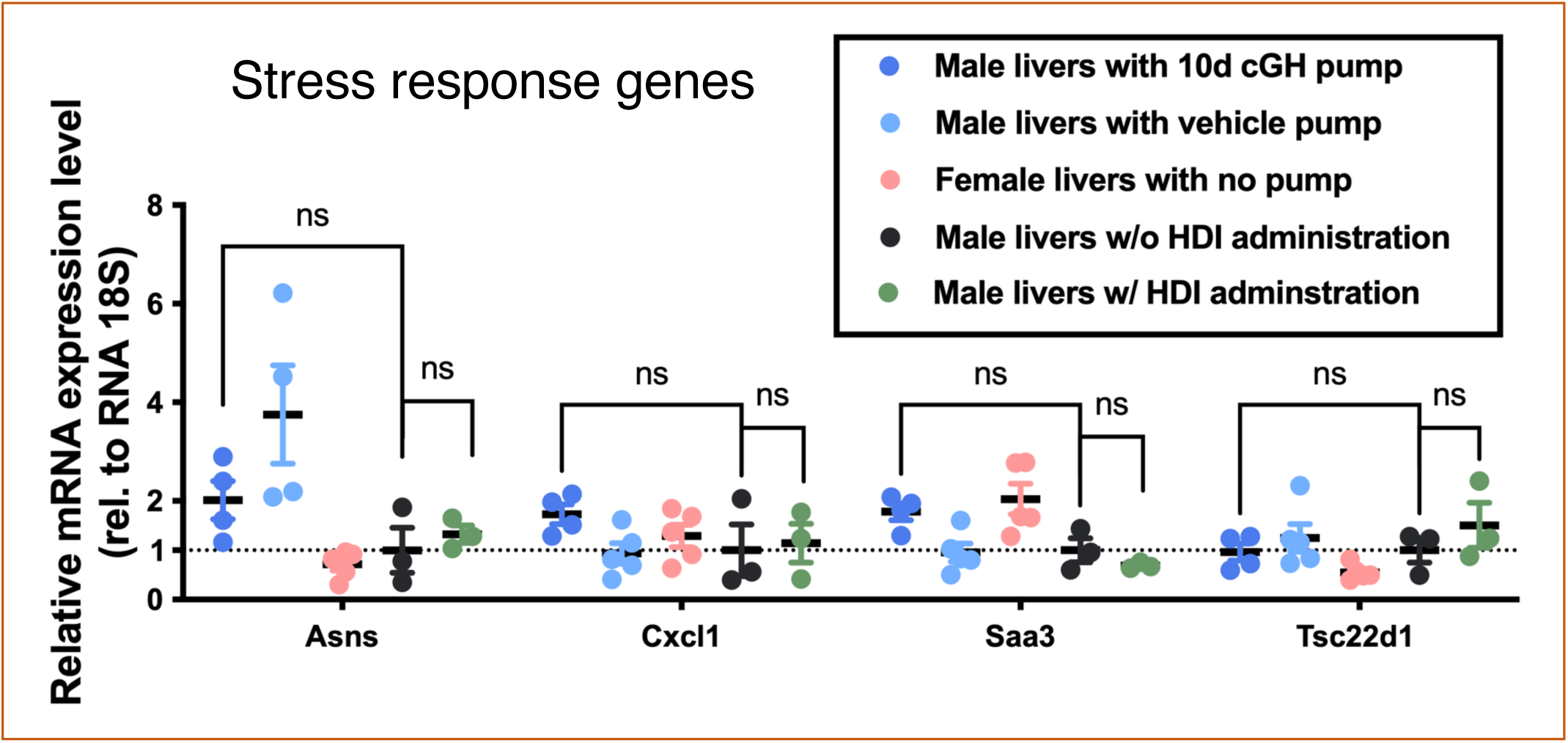
Stress gene expression. No stress genes were induced after 10 days of osmotic minipump implantation or 7 days after HDI. Statistical tests by Kolmogorov-Smirnov test (N=3-5 per group), followed by multiple comparison with FDR for multiple comparison correction. Data are mean +/− SEM values, with mean values for male without HDI set = 1.

**Fig. S8.**
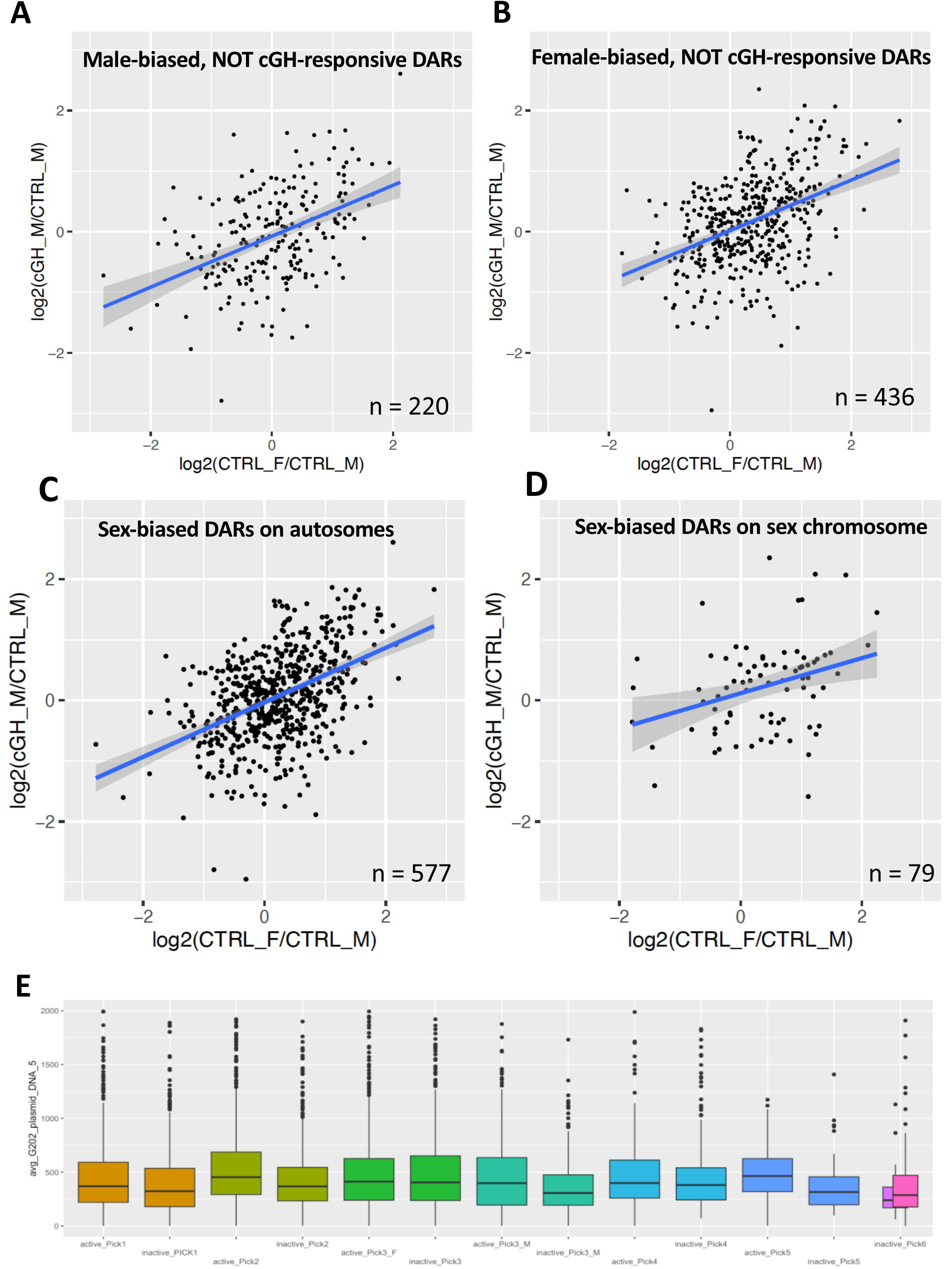
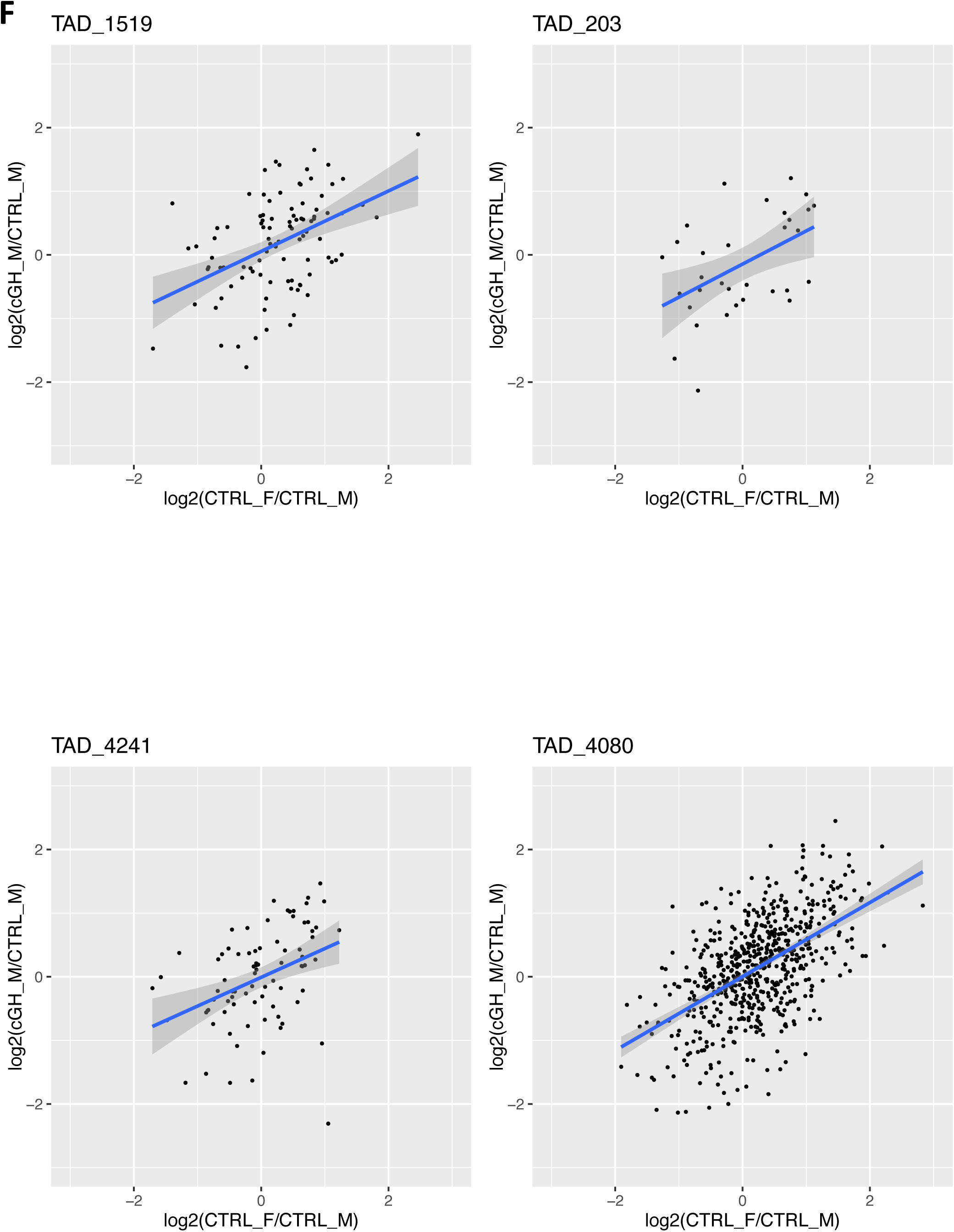
Correlation analysis of regulated sex-based and other reporters. **(A)** Correlation of cGH treatment and sex specificity in term of the ratio of STARR-seq reporter activity for the n=220 subset of 4,126 robust active reporters that are male-biased, not cGH-responsive DARs (Pick3_M, Table S2). **(B)** Correlation of cGH treatment and sex specificity in term of the ratio of STARR-seq reporter activity for the n=436 subset of 4,126 robust active reporters that are female-biased, not cGH-responsive DARs. Blue lines indicate linear regression models and gray region around each line represents the 95% confidence interval. **(C)** Correlation, as above, for n=577 autosomal robust active reporters that are at sex-biased DARs that are not cGH-responsive (Pick3_M and Pick3_F, Table S2). **(D)** Correlation as above for the n=79 robust active reporters on chrX or chrY that map to DARs that are sex-biased but not cGH-responsive. Blue lines in panels represent the best fit linear regression models, with the 95% confidence interval show in gray. **(E)** Boxplots indicate comparable plasmid library representation of the subsets of regulated and non-regulated reporters in each DAR set (‘Pick’ group, Table S2). **(F)** TAD-based correlations, as indicated. See Table S7B for detailed correlation data.

**Fig. S9.**
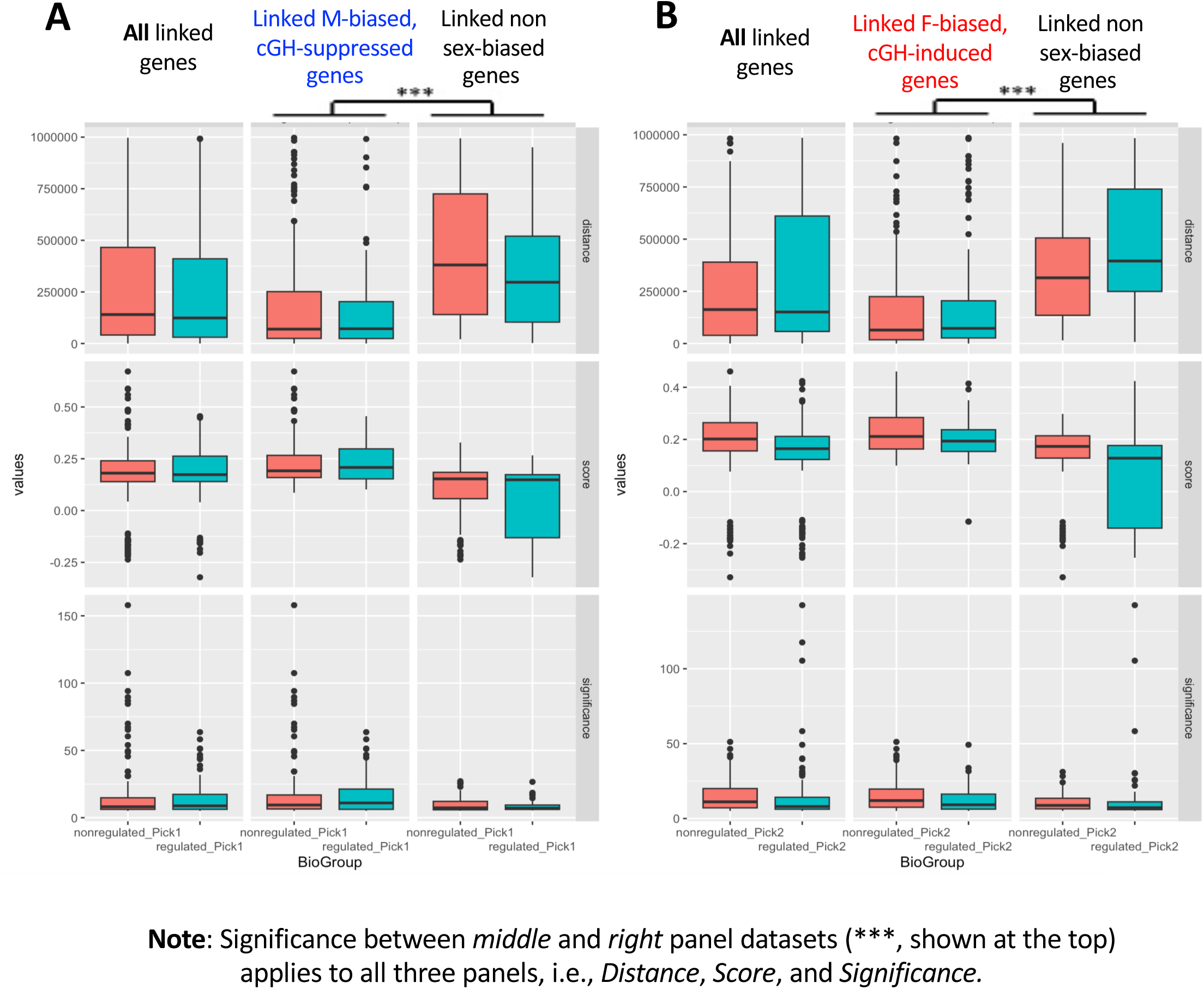
Gene linkage data for regulated vs non-regulated active enhancer DARs. Metrics of individual linkages between ATAC regions and genes grouped based on their HDI-STARR-seq activity and gene response, as derived from Table S4B and Table S9. Results show that linkages to non-sex-biased genes are characterized by significantly longer distances and significantly lower reliability (lower linkage score, lower linkage significance) than the linkages to sex biased genes; compare *middle* vs *right* subpanels of (A) and of (B). Shown are boxplots displaying the distributions of sex-biased regulated ATAC regions (salmon) or non-regulated regions (blue) with linkages to genes that are sex-biased [in (A), male biased and cGH-suppressed genes linked to regulated male-biased and cGH-suppressed DARs (i.e., 41 regulated regions); and in (B), female-biased and cGH-induced genes linked to regulated female-biased and cGH-activated DARs (i.e., 43 regulated regions)] in the *middle* panels, or that are linked to genes that are not sex-biased (*right* panels), or that are linked to any genes (*left* panels), where ‘any genes’ includes both sex-biased and not sex-biased genes (denoted as “all” in the subpanels). Statistical analyses were performed by applying the Kolmogorov-Smirnov test to the distributions for each of three metrics [i.e., linkage <u>distance</u> in nts (*top* subpanels), linkage <u>score</u> (*middle* subpanels), and linkage <u>significance</u> (−log(pvalue), *bottom* subpanels)] and are significant at p < 0.001 (***) for linkages to sex-biased genes versus linkage to non sex-biased genes, when considering the regulated regions and non-regulated regions as a group. In contrast, there are no significant differences (p-values >0.05, not marked in the figure) between regulated and non-regulated sex-biased and cGH-responsive DARs regarding the distributions of linkage scores, distances and significance, regardless of whether the linkage are to sex-biased genes or to non-sex-biased genes.

**Fig. S10,Fig. S11.**
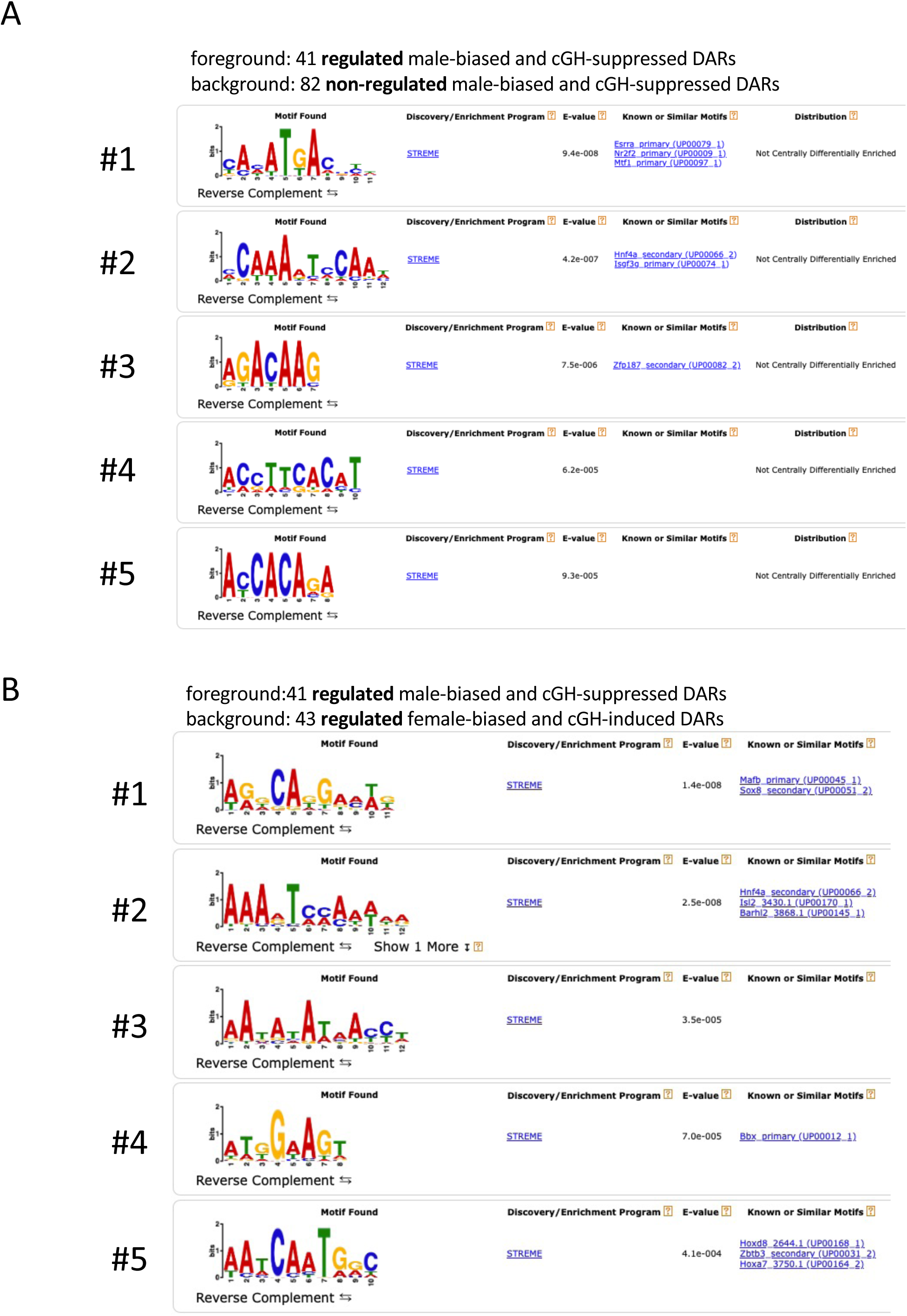
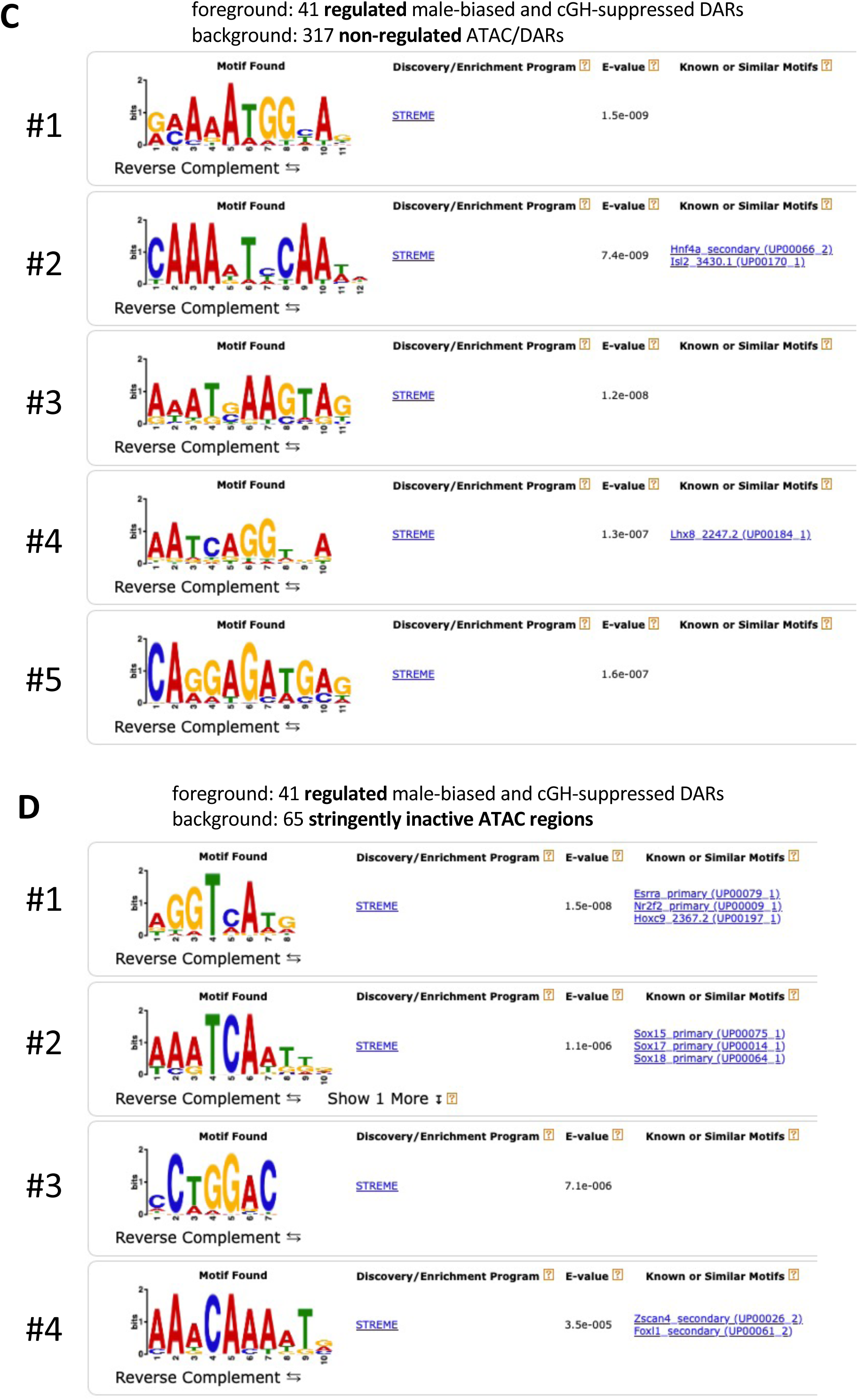
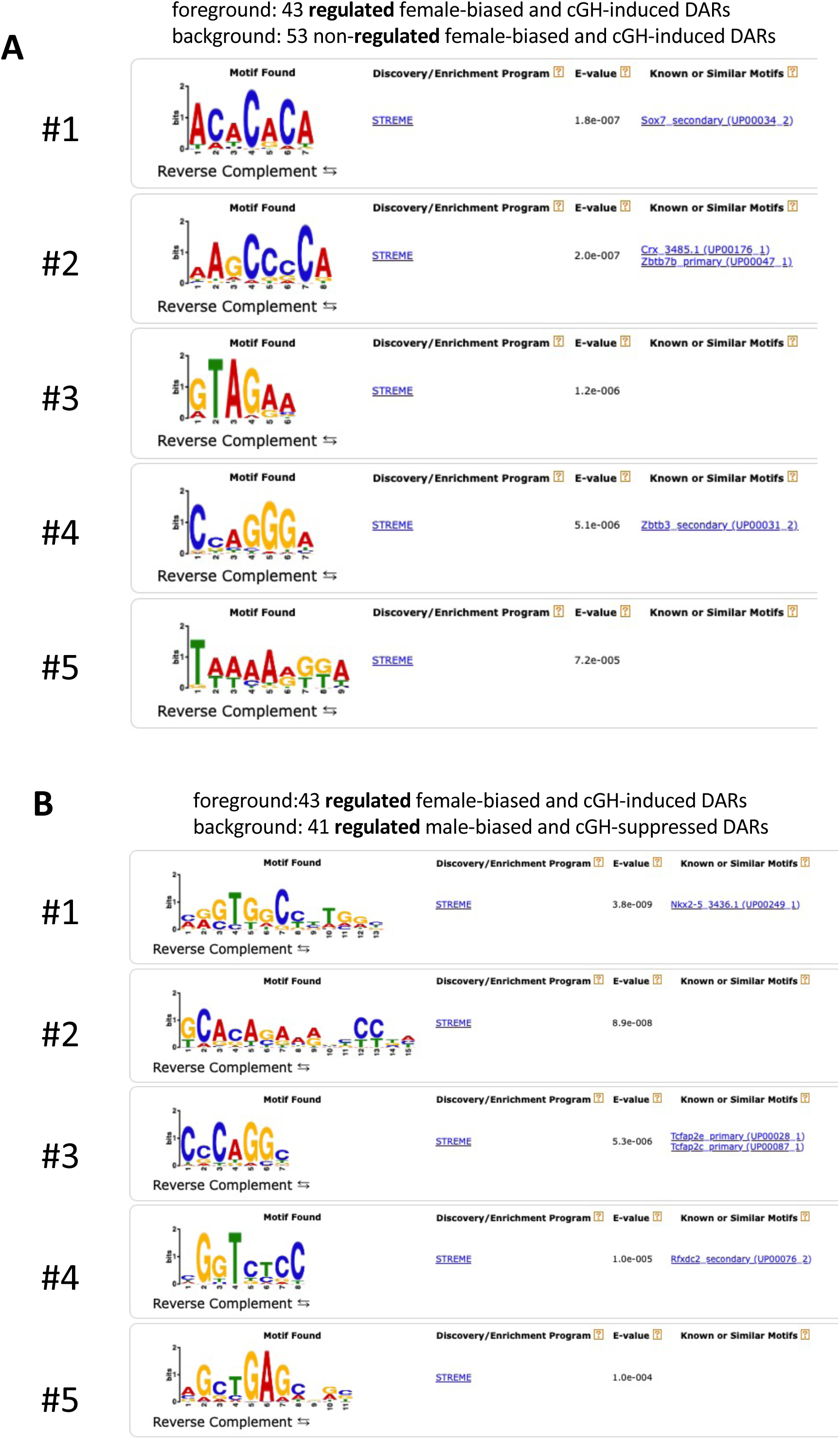
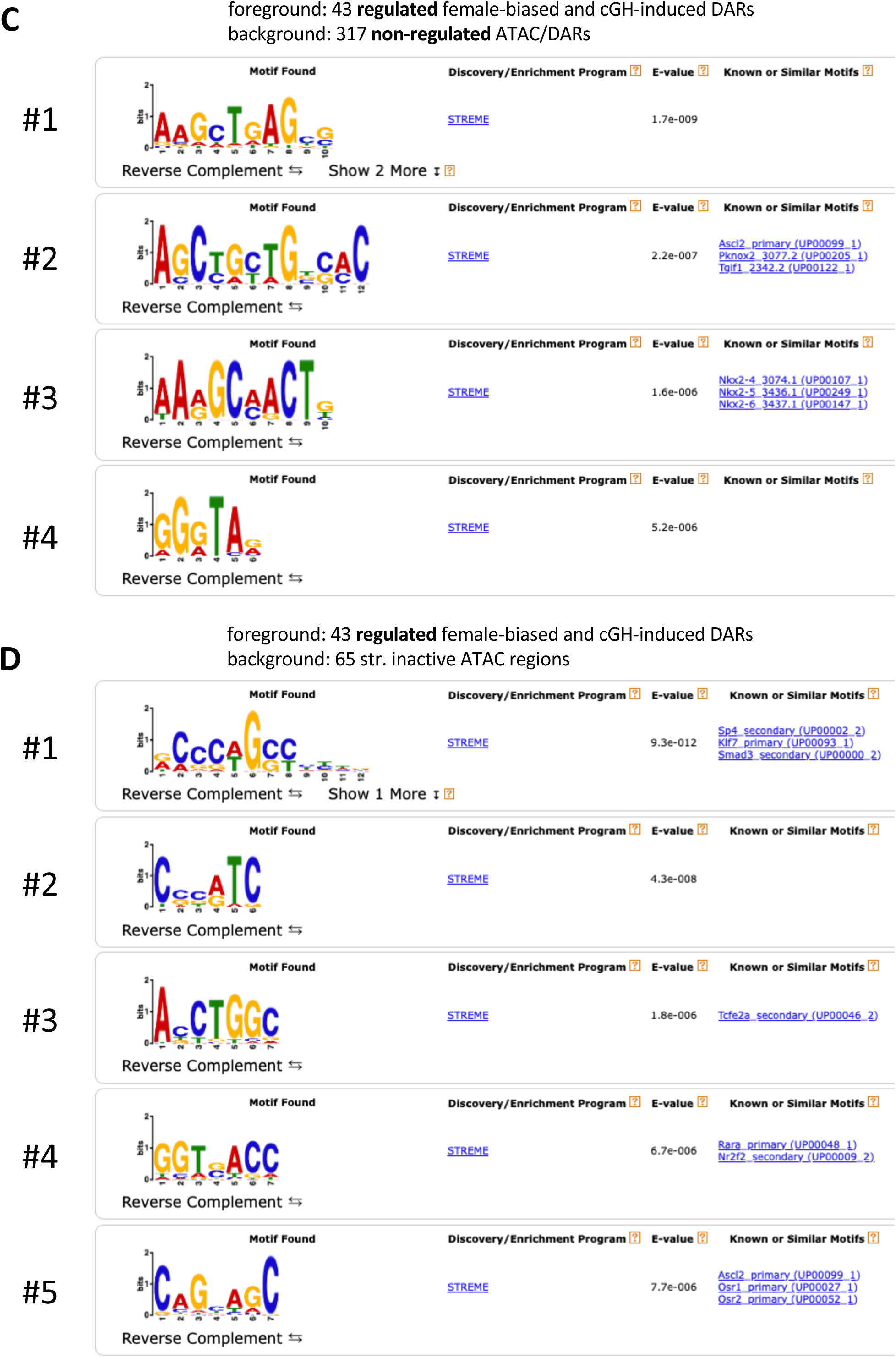
Motif analysis on regulated DARs. De novo motifs and the compared motifs in a mouse motif database (UniProbe) identified by MEME-ChIP for a variety of combination of foreground and background sets (see text).

**Fig. S12.**
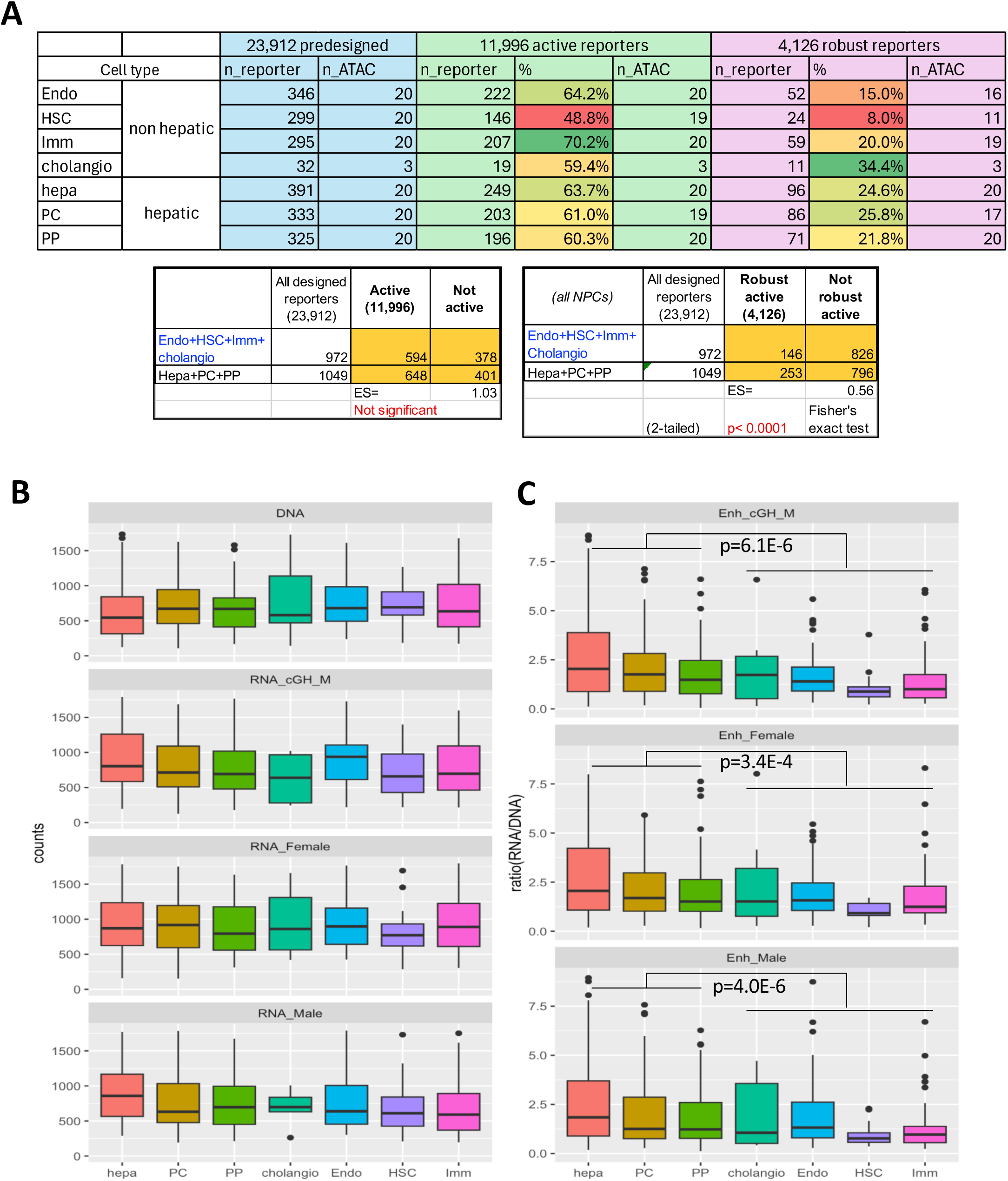
HDI-STARR-seq activity in cell-type specific ATAC regions. (A) Reporters representing liver cell-type specific ATAC regions included in the sets of 11,996 active reporters and 4,126 robust reporters identified in Table S7A. Percentages were calculated by dividing the number of active reporters (11,996 active reporters or 4,126 robust active reporters) by the 23,912 predesigned reporters in each cell type category. (A) Robust active reporters representing hepatic ATAC regions were significantly depleted in NPC ATAC regions. (B) Boxplots showing the distribution of DNA reads and RNA reads of the robust active reporters in each category. (C) Boxplots showing the distribution of enhancer activity for robust active reporters whose ATAC regions are preferentially accessible in the indicated liver cell type. Enhancer activity in hepatic ATAC regions was significantly higher in hepatocyte-specific ATAC regions than in NPC-specific ATAC regions (Kolmogorov-Smirnov test). Also see Table S11.

